# A shift in the mechanisms controlling hippocampal engram formation during brain maturation

**DOI:** 10.1101/2023.01.09.523283

**Authors:** Adam I. Ramsaran, Ying Wang, Ali Golbabaei, Bi-ru Amy Yeung, Mitchell L. de Snoo, Asim J. Rashid, Ankit Awasthi, Jocelyn Lau, Lina M. Tran, Sangyoon Y. Ko, Andrin Abegg, Lana Chunan Duan, Cory McKenzie, Julia Gallucci, Moriam Ahmed, Rahul Kaushik, Alexander Dityatev, Sheena A. Josselyn, Paul W. Frankland

## Abstract

The ability to form precise, episodic memories develops with age, with young children only able to form gist-like memories that lack precision. The cellular and molecular events in the developing hippocampus that underlie the emergence of precise, episodic-like memory formation are unclear. In mice, the absence of a competitive neuronal engram allocation process in the immature hippocampus precluded the formation of sparse engrams and precise memories until the fourth postnatal week, when inhibitory circuits in the hippocampus mature. This age-dependent shift in precision of episodic-like memories involved the functional maturation of parvalbumin-expressing interneurons in subfield CA1 by extracellular perineuronal nets which is necessary and sufficient for the onset of competitive neuronal allocation, sparse engram formation, and memory precision.

**One-Sentence Summary:** Episodic-like memory precision requires maturation of hippocampal inhibitory interneurons by the extracellular matrix.

---

The episodic memory system is absent or immature at birth and develops during childhood. Accordingly, early event memories are imprecise or gist-like until ∼5-8 years of age when mnemonic precision increases^1–5^. Hippocampal maturation is thought to underlie the emergence of precise episodic memories^5–8^, but the specific processes regulating memory precision during hippocampal development are unknown.

The binding of events to their surrounding spatial context is a core feature of episodic memory and may be studied in animals using spatial or contextual learning tasks^9^. To assess when this ability emerges during mouse development, we trained mice of different ages in contextual fear conditioning and tested their memory 24 hours later in either the same (context A) or a distinct (context B) testing apparatus (**Fig. 1A-B**). Younger mice (P16-P20) expressed imprecise contextual fear memories, freezing at equivalent levels in the training context A and the novel context B. In contrast, older mice (≥P24) expressed context-specific memories, freezing more in context A than context B (**Fig. 1C**). This shift in memory precision parallels similar shifts in rats^10^, was independent of the animals’ sex or weaning status and did not depend on prior experience with contexts, potential age-dependent differences in learning rate, or ability to perceptually discriminate the contexts (**Fig. S1A-Q**). Memory imprecision in juvenile mice scaled with the similarity between the training and testing contexts (**Fig. S1R-U).** Shifts in memory precision also occurred between P20 and P24 in a related aversive contextual learning task (inhibitory avoidance, **Fig. S1V-X**) and an appetitively-motivated spatial foraging task (**Fig. 1D-F, S2A-Q**).

**Fig. 1.**
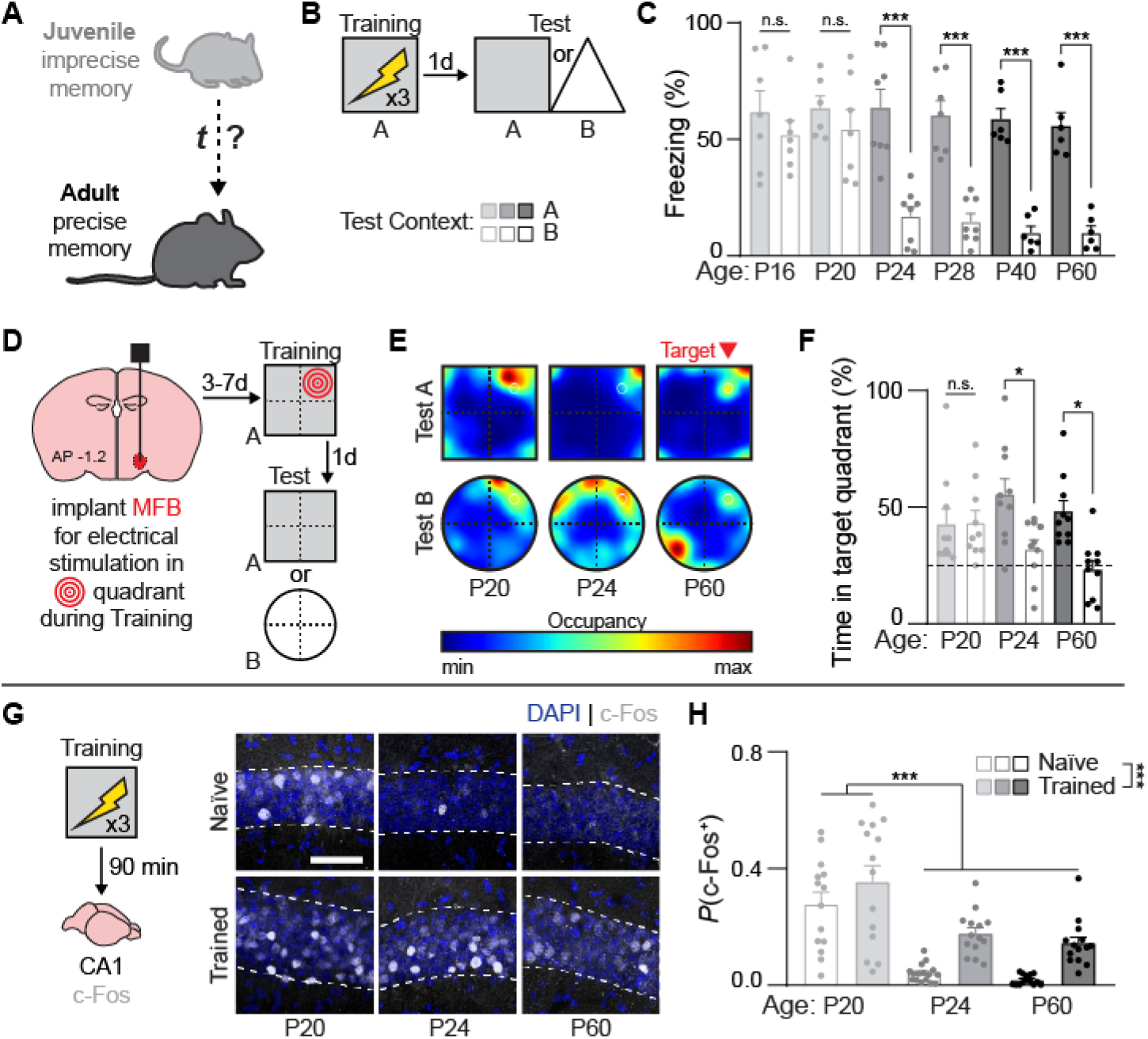
Memory precision and sparse engrams develop in dorsal CA1 during the fourth postnatal week. **(A)** The development of memory precision was assessed in mice. **(B)** Schematic of contextual fear conditioning protocol. **(C)** Juvenile mice (P16-P20) formed imprecise contextual fear memories, whereas older (>P24) mice formed precise memories (ANOVA, Age × Test Context interaction: *F*_1,70_ = 4.91, *P* < 0.001; main effect of Age: *F*_5,70_ = 6.74, *P* < 0.0001; main effect of Test Context: *F*_1,70_ = 92.34, *P* < 0.000001). **(D)** Schematic of spatial foraging task. **(E)** Heat maps depicting the average search pattern of P20, P24, and P60 mice during the test session. **(F)** Juvenile mice (P20) formed imprecise spatial memories, whereas older (>P24) mice formed precise memories (ANOVA, Age × Test Context interaction: *F*_2,54_ = 3.65, *P* < 0.05; no main effect of Age: *F*_2,54_ = 1.39, *P* = 0.25; main effect of Test Context: *F*_1,54_ = 13.43, *P* < 0.001). **(G)** c-Fos expression in dorsal CA1 90 min after contextual fear conditioning. Images show c-Fos expression in a segment of the dorsal CA1 pyramidal layer. **(H)** Approximately twice as many CA1 pyramidal layer cells expressed c-Fos after conditioning (or in home cage) in P20 mice compared to conditioned P24 and P60 mice (ANOVA, no Age × Experience interaction: *F*_2,80_ = 0.52, *P* = 0.59; main effect of Age: *F*_2,80_ = 34.81, *P* < 0.000001; main effect of Experience: *F*_1,80_ = 20.50, *P* < 0.0001). Data points are individual mice with mean ± s.e.m. Scale bars: white = 50 *μ*m, yellow = 500 *μ*m. * *P* < 0.05; ** *P* < 0.01; *** *P* < 0.001.

While contextual fear memories depend on the hippocampus in adult rodents^11^, it is possible that this type of learning is supported by extra-hippocampal structures in juvenile mice^12^. This is in line with proposals that the hippocampus does not support early event memories in children but instead ‘comes online’ during childhood to allow the emergence of episodic memory^12, 13^. We tested the hippocampal-dependency of contextual fear memories in juvenile and adult mice by microinjecting adeno-associated viruses (AAVs) encoding inhibitory opsins into dorsal CA1 of the hippocampus (**Fig. S3A-B**), as CA1 may support both precise and imprecise memories across development^14, 15^. Optogenetic silencing of CA1 pyramidal neurons impaired precise memory recall in P60 mice^11^ (**Fig. S3C-D**). In P20 mice, silencing CA1 neurons reduced freezing in both the A and B contexts., indicating that CA1 supports imprecise contextual memories at this developmental stage.

While these results indicate that the immature hippocampus supports early memories, age-dependent differences in memory precision suggest that these memories may be encoded differently in the hippocampi of juvenile vs. adult mice. In adults, context memories are encoded by sparse ensembles of neurons (also known as engrams) in the hippocampus^16, 17^. Given the imprecision of juvenile memories, we wondered whether CA1 engrams supporting contextual memories in juvenile mice lack sparsity. To identify putative engram neurons, we examined expression of the activity-regulated immediate-early gene (IEG) c-Fos in the dorsal CA1 of P20, P24, and P60 mice after contextual fear conditioning (**Fig. 1G**). Training induced Fos expression in ∼20% CA1 neurons in P24 and P60 mice^18–20^. In contrast, training induced c-Fos expression in ∼40% CA1 neurons in P20 mice, suggesting that engrams are more densely encoded in juvenile mice (**Fig. 1H**). Moreover, P20 mice showed a similar high proportion of c-Fos^+^ CA1 neurons in ‘home cage’ mice, consistent with observations that IEG expression is transiently elevated in the hippocampus of experimentally naive rodents during the third postnatal week as the activity-dependent assembly of hippocampal neural circuitry nears completion^21–23^ (**Fig. S4A-E**).

To test whether there is a causal relationship between engram size and memory precision, we asked whether artificially shrinking engrams in juvenile mice would promote adult-like memory precision. To sparsify the juvenile engram, we expressed the inhibitory DREADD (designer receptor exclusively activated by designer drugs) hM4Di in a subset of CA1 neurons and injected the DREADD ligand, C21, before training to inhibit infected neurons and prevent their inclusion in the engram (**Fig. 2A, S5A**). C21 treatment reduced engram size (training-induced c-Fos expression; **Fig. 2B-C**), and these juvenile mice precociously exhibited adult-like memory precision (freezing more in context A than context B) (**Fig. 2D-E, S5B-C)**.

**Fig. 2.**
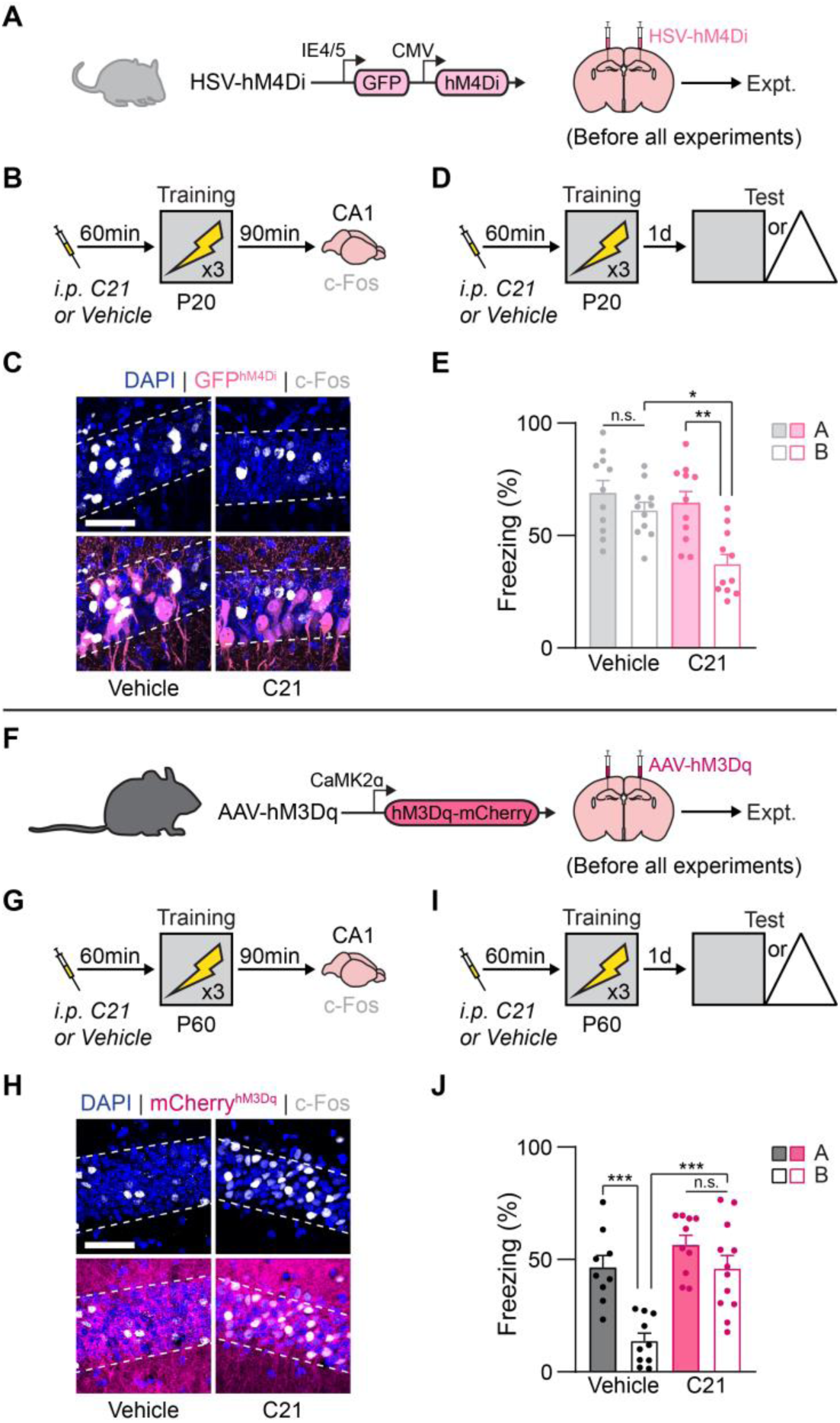
Engram sparsity in dorsal CA1 controls memory precision in juvenile and adult mice. **(A)** Engram size in CA1 of P20 mice was artificially decreased with HSV-hM4Di. **(B to C)** C21 was injected 1 h before training (B) and reduced c-Fos expression in the juvenile CA1 (C). **(D to E)** P20 mice administered C21 before training (D) formed precise contextual fear memories.Vehicle-treated P20 mice formed imprecise memories (E, ANOVA, Drug × Test Context interaction: *F*_1,26_ = 4.48, *P* < 0.05; no main effect of Drug: *F*_1,26_ = 3.02, *P* = 0.09; main effect of Test Context: *F*_1,26_ = 8.09, *P* < 0.01). **(F)** Engram size in CA1 of P60 mice was artificially increased with AAV-hM3Dq. **(G to H)** C21 was injected 1 h before training (G) and increased c-Fos expression in the adult CA1 (H) thereby promoting dense engram formation. **(I to J)** P60 mice administered C21 before training (I) formed imprecise contextual fear memories compared to vehicle-treated P60 mice that formed precise memories (J, ANOVA, Drug × Test Context interaction: *F*_1,37_ = 5.14, *P* < 0.05; main effect of Drug: *F*_1,37_ = 18.79, *P* < 0.001; main effect of Test Context: *F*_1,37_ = 19.62, *P* < 0.0001). Data points are individual mice with mean ± s.e.m. Scale bars: white = 50 *μ*m. * *P* < 0.05; ** *P* < 0.01; *** *P* < 0.001.

Conversely, we asked whether artificially expanding the engram in adult mice would induce juvenile-like memory imprecision. We expressed the excitatory DREADD construct, hM3Dq in pyramidal layer neurons (**Fig. 2F, S5D**) and injected C21 before training to increase the activity of hM3D-infected neurons. C21 treatment increased engram size (training-induced c-Fos expression; **Fig. 2G-H**), and these adult mice exhibited juvenile-like memory imprecision (equal freezing in contexts A and B) (**Fig. 2I-J, S5E-F)**. Thus, the delayed onset of adult-like memory functions by hyperactivity within developing memory circuits^24^ may be a core feature of ontogeny across animal species.

Such differences in engram sparsity between young and old mice suggest that the mechanisms of memory formation differ across development. In adult animals, eligible neurons are allocated to a sparse engram based on relative neuronal excitability or activity at the time of memory formation. To maintain engram sparsity, not only are neurons with relatively higher excitability included in the engram, but neurons with relatively lower excitability are excluded from the engram via lateral inhibition^25^. We probed how neuronal allocation changes across development using an all-optical strategy. We injected a replication-defective herpes simplex virus (HSV) into dorsal CA1 to infect a sparse random population of neurons with both a blue light-(BL) sensitive excitatory opsin (ChR2) and red light- (RL) sensitive inhibitory opsin (eNpHR3.0) (HSV-NpACY, **Fig. 3A-B, S6A**). This approach allowed us to bidirectionally modulate the activity of the same population of infected neurons with different wavelengths of light. Similar to previous experiments in the lateral amygdala^26^, we briefly excited NpACY^+^ neurons with BL immediately before conditioning to bias their allocation into the engram. Control mice were treated similarly but received no BL. Following training, c-Fos was preferentially expressed in NpACY^+^ neurons in P20, P24 and P60 mice in the BL+ (Allocated) but not BL- (Control, Non-allocated) mice (**Fig. S6B-E**), suggesting that optogenetic-mediated allocation was effective regardless of mouse age.

**Fig. 3.**
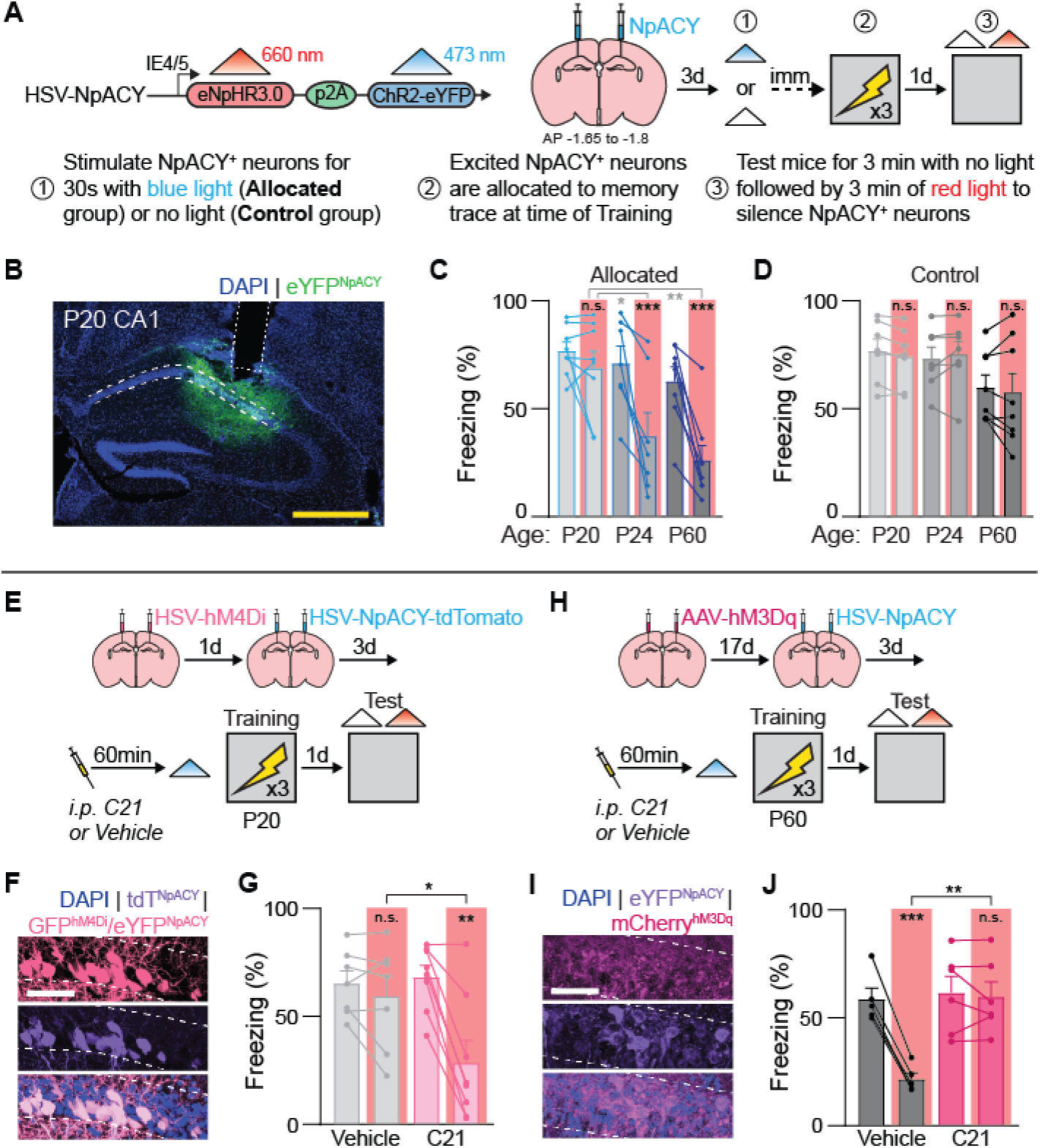
Immature neuronal allocation mechanisms in CA1 preclude localization of memories to sparse engrams during early development. **(A)** Schematic of the ‘allocate-and-silence’ contextual fear conditioning protocol. HSV-NpACY was used to optogenetically excite (ChR2) and inhibit (NpHR3.0) the same neurons. **(B)** HSV-NpACY expression in dorsal CA1 of a P20 mouse. **(C)** Silencing a sparse group of NpACY^+^ neurons previously allocated to the contextual fear memory engram impaired freezing during the test in P24 and P60 mice, but not P20 mice (RM-ANOVA, Age × Light interaction: *F*_2,20_ = 4.96, *P* < 0.05; main effect of Age: *F*_2,20_ = 5.09, *P* < 0.05; main effect of Light: *F*_1,20_ = 39.28, *P* < 0.00001). **(D)** Silencing a sparse group of random NpACY^+^ neurons not allocated to the engram did not impair freezing during the test in any mice (RM-ANOVA, Age × Light interaction: *F*_2,19_ = 1.05, *P* = 0.36; no main effect of Age: *F*_2,19_ = 2.38, *P* = 0.11; no main effect of Light: *F*1,19 = 0.45, *P* = 0.50). **(E)** HSV-hM4Di and HSV-NpACY were both expressed in CA1 before contextual fear conditioning. C21 was administered 1 h before training to promote sparse engram formation, and HSV-NpACY was used to allocate a sparse group of neurons to the engram at the time of training and silence the same neurons during the test. **(F)** Expression of NpACY and hM4Di in dorsal CA1 of a P20 mouse. **(G)** Silencing a sparse group of NpACY^+^ neurons previously allocated to the contextual fear memory engram impaired freezing during the test in P20 mice that received C21 to shrink their engrams (RM-ANOVA, Drug × Light interaction: *F*_1,13_ = 14.25, *P* < 0.01; no main effect of Drug: *F*_1,13_ = 1.79, *P* = 0.20; main effect of Light: *F*_1,13_ = 25.71, *P* < 0.001). **(H)** AAV-hM3Dq and HSV-NpACY were both expressed in CA1 before contextual fear conditioning. C21 was administered 1 h before training to promote dense engram formation, and HSV-NpACY was used to allocate a sparse group of neurons to the engram at the time of training and silence the same neurons during the test. **(I)** Expression of NpACY and hM3Dq in dorsal CA1 of a P60 mouse. **(J)** Silencing a sparse group of NpACY^+^ neurons previously allocated to the contextual fear memory engram did not impair freezing during the test in P60 mice that received C21 to expand their engrams (RM-ANOVA, Drug × Light interaction: *F*_1,9_ = 63.22, *P* < 0.0001; main effect of Drug: *F*_1,9_ = 5.67, *P* < 0.05; main effect of Light: *F*_1,9_ = 77.56, *P* < 0.0001).

In a second cohort of mice, we repeated the same allocation procedure and then probed whether infected neurons were necessary for subsequent memory expression by using RL to silence NpACY^+^ expressing neurons during a memory test. Silencing NpACY^+^ neurons impaired fear recall in P24 and P60 mice in the Allocated (but not Control, Non-allocated) groups (**Fig. 3C-D**), indicating that the CA1 engram was localized to the sparse NpACY^+^ population of neurons. In contrast, silencing a similar number of NpACY^+^ neurons did not impair fear memory recall in P20 mice. This suggests that information is more broadly distributed in densely-encoded engrams in juvenile mice, such that silencing only a fraction of these neurons is not sufficient to disrupt memory recall. The dense juvenile engram included both artificially-allocated (NpACY^+^c-Fos^+^) and non-allocated (NpACY^-^c-Fos^+^) neurons (**Fig. S6F**).

In juvenile mice, chemogenetic shrinking of the engram promoted adult-like neuronal allocation. Silencing allocated NpACY^+^ neurons impaired fear recall in the juvenile C21 group (**Fig. 3E-G**), indicating that the CA1 engram was localized to the sparse NpACY^+^ population of neurons in P20 mice with artificially-shrunken engrams. In adult mice, chemogenetic expansion of the engram induced juvenile-like neuronal allocation. Silencing allocated NpACY^+^ neurons no longer impaired fear memory recall in the adult C21 group (**Fig. 3H-J**), consistent with the idea that information was not localized to the sparse population of NpACY^+^ neurons, but more broadly distributed within the artificially-expanded engram.

The denser engrams in P20 mice raises the possibility that the second component of the mature neuronal allocation process — the exclusion of less-active or inactive neurons from engrams by interneurons — is not yet fully developed in juvenile mice. In the lateral amygdala, parvalbumin-expressing (PV^+^) basket cells provide strong somatic inhibition onto excitatory neurons at the time of memory formation to exclude less excitable neurons from the engram^26, 27^. In the CA1, PV^+^ interneurons are born embryonically in the medial ganglionic eminence, resulting in adult-like levels of these cells by the third postnatal week in mice^6, 28^ (**Fig. 4A-B**). Despite their early birthdate, morphological and functional development of CA1 PV^+^ interneurons continues into the fourth postnatal week in mice^29, 30^. Therefore, in P20 mice, developing PV^+^ interneurons in the CA1 may provide only weak lateral inhibition, precluding the formation of sparse engrams at the time of memory encoding. Adult-like levels of PV^+^ neurites and Syt2^+^ puncta (labeling PV^+^ interneuron presynaptic terminals, **Fig. S7A**) only emerged in the CA1 pyramidal layer at P24 (**Fig. 4C-D**). Moreover, contextual fear conditioning increased perisomatic PV staining surrounding c-Fos^-^ (cf. c-Fos^+^) pyramidal layer cells at P24 and P60, but not P20, consistent with the idea that experience-dependent lateral inhibition is not occurring until the fourth postnatal week (**Fig. 4E-F, S7B-C**).

**Fig. 4.**
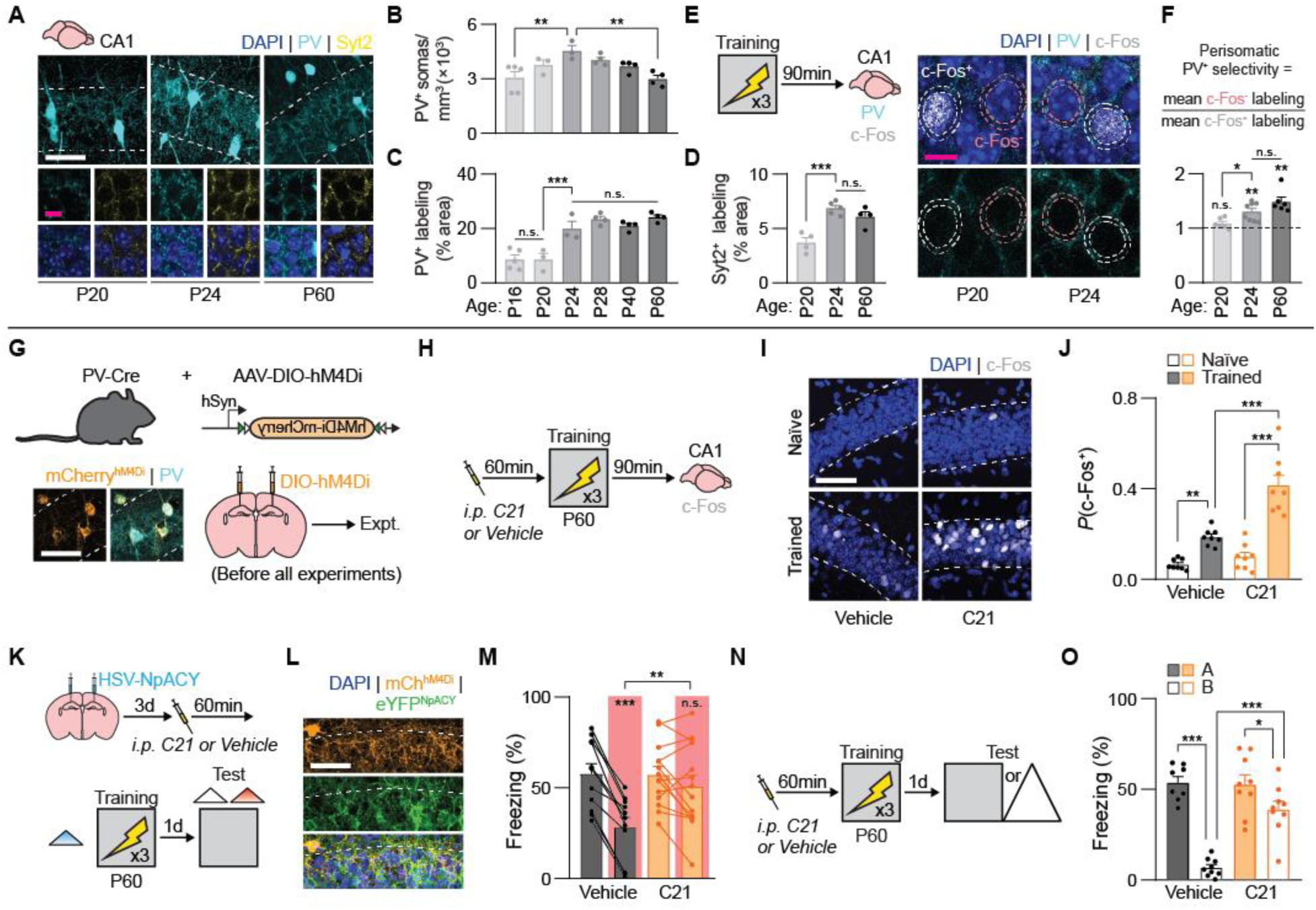
Mature CA1 PV^+^ interneuron function is required for competitive neuronal allocation, sparse engram formation, and memory precision. **(A)** PV^+^ interneurons, PV^+^ neurites, and Syt2^+^ terminals in CA1 across development. **(B)** The number of PV^+^ interneurons peaks transiently at P24 (ANOVA, effect of Age: *F*_5,17_ = 5.02, *P* < 0.01). **(C)** The density of PV^+^ neurites in the pyramidal layer reaches adult-like levels by P24. (ANOVA, effect of Age: *F*_5,17_ = 21.78, *P* < 0.00001). **(D)** The density of Syt2^+^ synaptic terminals in the pyramidal layer reaches adult-like levels by P24. (ANOVA, effect of Age: *F*_2,10_ = 18.01, *P* < 0.001). **(E)** c-Fos and PV expression were examined in dorsal CA1 90-min after contextual fear conditioning. Localization of PV^+^ neurites around c-Fos^-^ and c-Fos^+^ pyramidal layer cells. PV^+^ labeling was quantified in a 3-μm ring surrounding the nuclei. **(F)** PV^+^ neurites were selectively localized (Perisomatic PV^+^ Selectivity > 1) around c-Fos^-^ compared to c-Fos^+^ cells after training in P24 and P60 mice, but not P20 mice (one-sample *t*-tests [with Bonferroni correction, ɑ = 0.016], P20: *t*_5_ = 2.25, *P* = 0.073; P24: *t*_6_ = 5.16, *P* < 0.01; P60: *t*_5_ = 5.85, *P* < 0.01; ANOVA, effect of Age: *F*_2,17_ = 9.04, *P* < 0.01). **(G)** PV^+^ interneurons were inhibited by expressing AAV-DIO-hM4Di in CA1 of adult PV-Cr mice. **(H)** c-Fos expression in dorsal CA1 was examined 90-min after contextual fear conditioning. C21 was administered 1 h before training to inhibit PV^+^ interneurons. **(I)** c-Fos expression in the dorsal CA1 pyramidal layer. **(J)** Inhibiting PV^+^ interneurons before training resulted in a two-fold increase in c-Fos expression after training (ANOVA, Drug × Experience interaction: *F*_1,28_ = 13.82, *P* < 0.001; main effect of Drug: *F*_1,28_ = 24.67, *P* < 0.0001; main effect of Experience: *F*_1,28_ = 69.14, *P* < 0.000001). **(K)** Schematic of the contextual fear conditioning protocol. C21 was administered 1 h before training to inhibit PV^+^ interneurons, and HSV-NpACY was used to excite (ChR2) and inhibit (NpHR3.0) the same neurons. **(L)** Expression of NpACY and hM4Di in dorsal CA1 of P6 mouse. **(M)** Silencing a sparse group of NpACY^+^ neurons previously allocated to the contextual fear memory engram did not impair freezing during the test in P60 mice that received C21 to inhibit their PV^+^ interneurons (RM-ANOVA, Drug × Light interaction: *F*_1,23_ = 21.12, *P* < 0.001; no main effect of Drug: *F*_1,23_ = 2.21, *P* = 0.14; main effect of Light: *F*_1,23_ = 51.61, *P* < 0.000001). **(N)** Schematic of the contextual fear conditioning protocol. **(O)** P60 mice administered C21 to inhibit PV^+^ interneurons formed imprecise contextual fear memories.Vehicle-treated P20 mice formed precise memories (ANOVA, Drug × Test Context interaction: *F*_1,31_ = 16.56, *P* < 0.001; main effect of Drug: *F*_1,31_ = 14.65, *P* < 0.001; main effect of Test Context: *F*_1,31_ = 55.73, *P* < 0.00001). Data points are individual mice with mean ± s.e.m. Scale bars: magenta = 10 *μ*m, whit = 50 *μ*m. * *P* < 0.05; ** *P* < 0.01; *** *P* < 0.001.

We next inhibited PV^+^ interneurons in CA1 by expressing hM4Di in PV-Cre, adult mice (**Fig. 4G, S8A-B**). Inhibition of PV^+^ interneurons with C21 before fear conditioning disinhibited local excitatory neurons, resulting in dense engrams that induced high levels of c-Fos in the CA1 after training (**Fig. 4H-J**). Moreover, inhibiting CA1 PV^+^ interneurons in adult mice promoted juvenile-like allocation and memory imprecision (**Fig. 4K-O, S8C-D**). Reinstatement of these juvenile-like mnemonic phenotypes in adult PV-Cre mice required the presence of both hM4Di and C21 at the time of memory encoding (and not memory retrieval) (**Fig. S8E-H**). Thus, acute PV^+^ interneuron inhibition at the time of memory formation was sufficient to phenocopy the allocation and memory specificity profile of juvenile mice, while also maintaining the critical role of CA1 in encoding and retrieving these modified memories (**Fig. S8I-L**). These experiments identify maturation of CA1 inhibitory PV^+^ circuitry as the key driver of memory development during the fourth postnatal week.

PV^+^ interneurons play a pivotal role in the development of cortical sensory systems by closing transient windows of critical period plasticity^31^. For example, PV^+^ interneuron maturation closes the critical period for ocular dominance plasticity in the binocular primary visual cortex (V1b) in mice and rats^32^. The maturation of perineuronal nets (PNNs), extracellular matrix (ECM) structures primarily ensheathing the soma and proximal dendrites of PV^+^ interneurons, helps drive PV^+^ cell maturation in the cortex and hippocampus. By stabilizing excitatory synapses onto PV^+^ interneurons and inhibitory synapses originating from PV^+^ interneurons, mature PNNs increase PV^+^ interneuron-mediated inhibition^33–35^. Because PNN formation in V1b is necessary for the emergence of adult-like visual acuity, we reasoned that PNN formation in the hippocampus may similarly regulate the development of adult-like mnemonic specificity.

Mature PNNs represent a form of dense ECM composed of polymer chains of hyaluronan, chondroitin sulfate proteoglycans (CSPGs), tenascin-R, and link proteins^36, 37^. To establish a detailed developmental profile of PNNs in the hippocampus, we visualized *Wisteria floribunda* agglutinin (WFA), CSPGs, and link proteins including CSPG brevican (BCAN) and hyaluronan and proteoglycan link protein 1 (HAPLN1) *in situ*. WFA^+^ PNNs were found throughout the adult hippocampus (**Fig. S9A-C**). Hippocampal PNNs primarily surrounded PV^+^ interneurons (with the exception of fasciola cinereum and CA2 PNNs) (**Fig. S9D-I**) and mostly developed postnatally. In CA1, adult-like levels of PNNs surrounding PV^+^ interneurons were achieved by P24 in both male and female mice (**Fig. 5A-D, S10A-F**). CA1 PNN density was not altered by contextual fear conditioning (**Fig. S10Q-R**), suggesting that only micro-scale PNN alterations are induced by memory formation^38^. In contrast to CA1, PV interneuron-associated PNNs reached adult levels before P16 in the DG and CA3 (**Fig. S11A-K**), consistent with the step-wise maturation of hippocampal trisynaptic circuitry^21^.

**Fig. 5.**
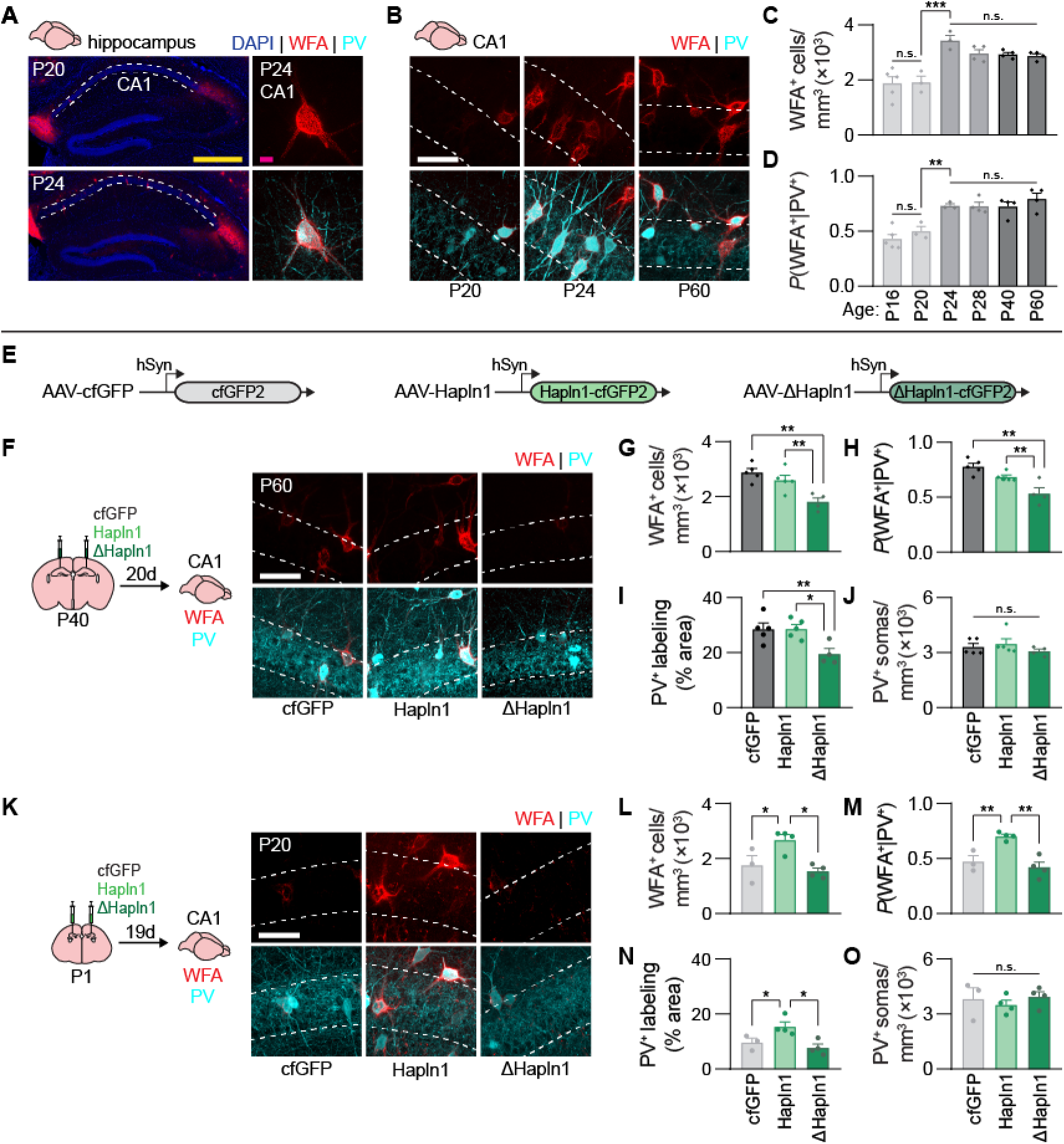
Maturation of CA1 PNNs and PV^+^ interneurons requires HAPLN1. **(A)** WFA^+^ PNNs in the hippocampus of a P20 and P24 mouse. High magnification image of a WFA^+^ PNN on P24 surrounding the soma and proximal dendrites of a PV^+^ interneuron. **(B)** WFA^+^ PNNs surrounding PV^+^ interneurons in CA1 across development. **(C to D)** The density of WFA^+^ PNNs in CA1 (ANOVA, effect of Age: *F*_5,17_ = 11.88, *P* < 0.0001) surrounding PV^+^ interneurons (ANOVA, effect of Age: *F*_5,17_ = 13.51, *P* < 0.0001) reached adult-like levels by P24. **(E)** To bidirectionally manipulate PNN integrity *in vivo*, we used AAV-Hapln1, AAV-ΔHapln1, and AAV-cfGFP to overexpress wild-type mouse HAPLN1, or a mutated dominant-negative ΔHAPLN1, or cysteine-free GFP as a negative control, respectively. **(F)** Mice were microinjected with AAVs and P60 brains were stained for WFA^+^ PNNs and PV^+^ interneurons in dorsal CA1. **(G to J)** Expression of AAV-ΔHapln1 in P60 CA1 decreased the number of WFA^+^ PNNs (G, ANOVA, effect of Virus: *F*_2,11_ = 10.88, *P* < 0.01) surrounding PV^+^ interneurons (H, ANOVA, effect of Virus: *F*_2,11_ = 12.52, *P* < 0.01) and density of PV^+^ neurites in the pyramidal layer (I, ANOVA, effect of Virus: *F*_2,11_ = 6.88, *P* < 0.05), without affecting the number of PV^+^ interneurons (J, ANOVA, no effect of Virus *F*_2,11_ = 0.89, *P* = 0.43). **(K)** Mice were microinjected with AAVs and P20 brains stained for WFA^+^ PNNs and PV^+^ interneurons in dorsal CA1. **(L to O)** Expression of AAV-Hapln1 in P20 CA1 increased the number of WFA^+^ PNNs (L, ANOVA, effect of Virus: *F*_2,8_ = 8.02, *P* < 0.05) surrounding PV^+^ interneurons (M, ANOVA, effect of Virus: *F*_2,8_ = 14.43, *P* < 0.01) and density of PV^+^ neurites in the pyramidal layer (N, ANOVA, effect of Virus: *F*_2,8_ = 7.47, *P* < 0.05), without affecting the number of PV^+^ interneurons (O, ANOVA, no effect of Virus: *F*_2,8_ = 0.42, *P* = 0.66). Data points are individual mice with mean ± s.e.m. Scale bars: magenta = 10 *μ*m, white = 50 *μ*m, yellow = 500 *μ*m. * *P* < 0.05; ** *P* < 0.01; *** *P* < 0.001.

Next, we tested whether PNN maturation controls memory development by manipulating PNN integrity in the CA1 of adult and juvenile mice. HAPLN1 plays a key role in cross-linking and stabilizing CSPG-hyaluronan interactions, and *Hapln1* transcription corresponds with mature PNN formation in the cortex^39^. To promote or interfere with PNN integrity, we designed viral vectors to overexpress wild-type (AAV-Hapln1), mutant (AAV-ΔHapln1) dominant-negative HAPLN1 proteins (not binding to CSPGs^40^) tagged with cfGFP or AAV-cfGFP control (**Fig. 5E**). Expression of the HAPLN1-targeting constructs did not affect total endogenous expression of PNN proteins in the dorsal hippocampus of adult mice (**Fig. S12A-L**). However, AAV-ΔHapln1 expression diminished CA1 PNN growth in P60 mice and consequently reduced the density of PV^+^ neurites in the CA1 pyramidal layer (**Fig. 5F-J**). In contrast, AAV-Hapln1 expression accelerated CA1 PNN growth and increased the density of PV^+^ neurites in the CA1 pyramidal layer in P20 mice (**Fig. 5K-O**).

First, we tested whether destabilizing CA1 PNNs in adult mice with AAV-ΔHapln1 would phenocopy the formation of a dense engram and memory imprecision typical of juvenile mice (**Fig. 6A-B**). Adult mice expressing AAV-ΔHapln1 (but not control protein or wild-type Hapln1) before training formed a dense engram (high proportion of c-Fos^+^ neurons following contextual fear conditioning) and intact memory when the sparse population of artificially allocated (HSV-NpACY-expressing) neurons were silenced during testing (**Fig. 6C-G**). PNN destabilization in separate groups of adult mice produced imprecise contextual fear (**Fig. 6H-I, S13A-B**) and appetitive spatial (**Fig. 6J-L, S14A-K**) memories. The effects of ΔHapln1 in adult mice were specific to episodic-like memory formation. Similar destabilization of CA1 PNNs with this construct did not decrease precision for auditory fear memories or increase anxiety-like behavior or locomotion in an open field (**Fig. S13C-G)**. These changes in contextual memory precision were largely restricted to HAPLN1-based manipulation of PNNs in CA1 (and not cortex) (**Fig. S13H-K**). We observed similar behavioral effects using the enzyme ChABC^35^ to transiently digest CA1 PNNs (**Fig S15A-R**).

**Fig. 6.**
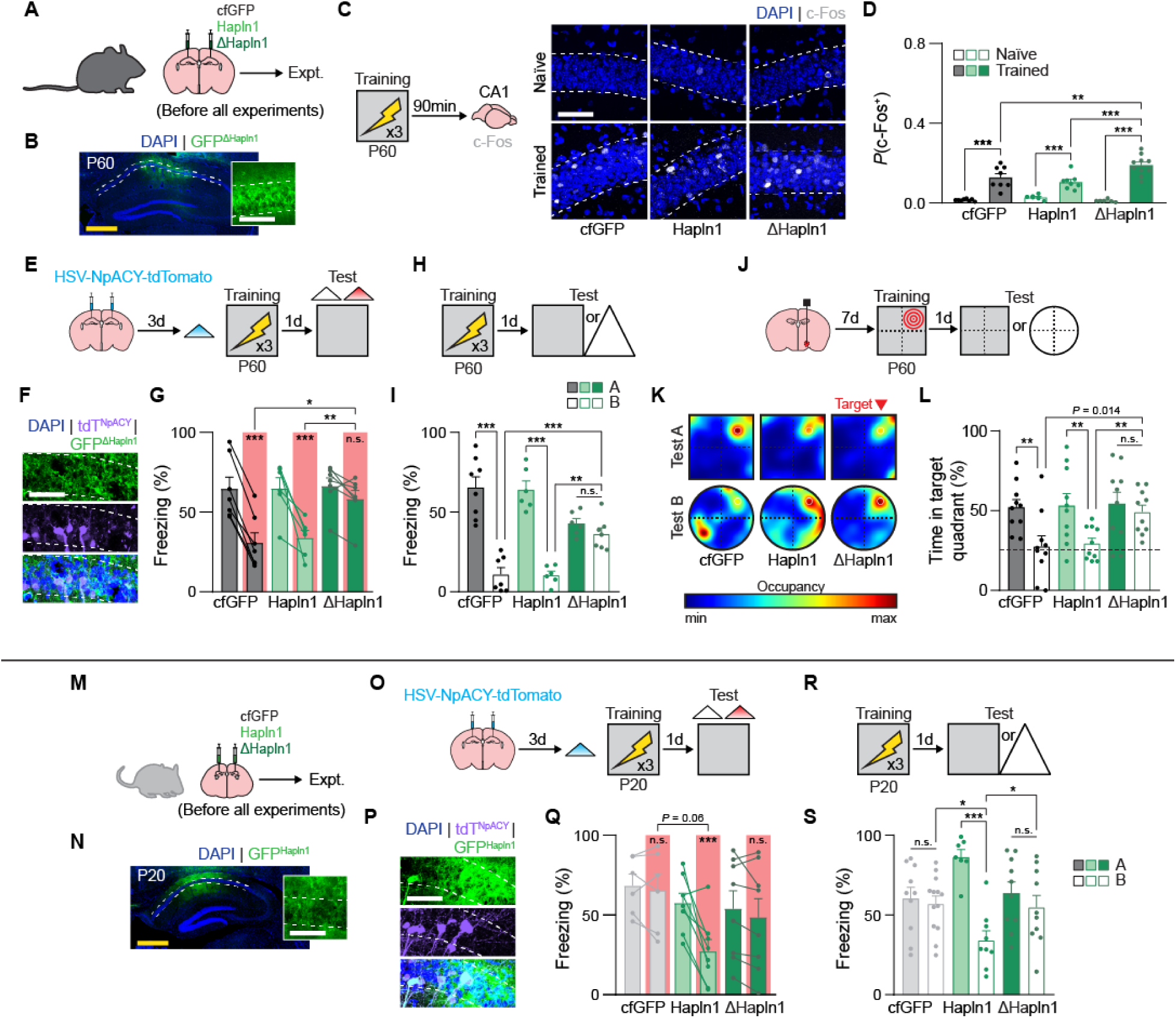
Maturation of CA1 PNNs is required for competitive neuronal allocation, sparse engram formation, and memory precision. **(A)** PNNs were manipulated in the adult CA1 by microinjecting AAVs before each experiment. **(B)** AAV-ΔHapln1 expression in the P60 CA1. **(C)** c-Fos expression in dorsal CA1 was examined 90-min after contextual fear conditioning. **(D)** Expression of AAV-ΔHapln1 in P60 CA1 before training resulted in a two-fold increase in c-Fos expression after training (ANOVA, Virus × Experience interaction: *F*_2,40_ = 6.89, *P* < 0.01; main effect of Virus: *F*_2,40_ = 3.44, *P* < 0.05; main effect of Experience: *F*_1,40_ = 122.83, *P* < 0.000001). **(E)** Schematic of the contextual fear conditioning protocol. HSV-NpACY was used to optogenetically excite (ChR2) and inhibit (NpHR3.0) the same neurons. **(F)** Expression of NpACY and AAV-ΔHapln1 in dorsal CA1 of P60 mouse. **(G)** Silencing a sparse group of NpACY^+^ neurons previously allocated to the contextual fear memory engram did not impair freezing during the test in P60 mice expressing AAV-ΔHapln1 in CA1 (RM-ANOVA, Virus × Light interaction: *F*_2,17_ = 9.58, *P* < 0.01; no main effect of Virus: *F*_2,17_ = 2.16, *P* = 0.14; main effect of Light: *F*_1,17_ = 82.46, *P* < 0.000001). **(H)** Schematic of the contextual fear conditioning protocol. **(I)** P60 mice expressing AAV-ΔHapln1 in CA1 formed imprecise contextual fear memories. P60 mice expressing AAV-cfGFP or AAV-Hapln1 formed precise memories (ANOVA, Virus × Test Context interaction:*F*_2,34_ = 16.40, *P* < 0.0001; no main effect of Virus: *F*_2,34_ = 0.11, *P* = 0.89; main effect of Test Context: *F*_1,34_ = 96.02, *P* < 0.00001). **(J)** Schematic of the spatial foraging task. **(K)** Heat maps depicting example search patterns during the test. **(L)** P60 mice expressing AAV-ΔHapln1 in CA1 formed imprecise spatial memories. P60 mice expressing AAV-cfGFP or AAV-Hapln1 formed precise memories (unpaired *t*-tests [with Bonferroni correction, ɑ = 0.01], AAV-cfGFP A vs. B: *t*_18_ = 2.99, *P* < .01; AAV-Hapln1 A vs. B: *t*_18_ = 2.90, *P* < .01; AAV-ΔHapln1 A vs. B: *t*_18_ = 0.59, *P* = 0.56; AAV-cfGFP B vs. AAV-ΔHapln1 B: *t*_18_ = 2.70, *P* = 0.014; AAV-Hapln1 B vs. AAV-ΔHapln1 B: *t*_18_ = 3.61, *P* < 0.01). **(M)** PNNs were manipulated in the juvenile CA1 by microinjecting AAVs before each experiment. **(N)** AAV-Hapln1 expression in the P20 CA1. **(O)** Schematic of the contextual fear conditioning protocol. HSV-NpACY was used to optogenetically excite (ChR2) and inhibit (NpHR3.0) the same neurons. **(P)** Expression of NpACY and AAV-Hapln1 in dorsal CA1 of P20 mouse. **(Q)** Silencing a sparse group of NpACY^+^ neurons previously allocated to the contextual fear memory engram impaired freezing during the test in P20 mice expressing AAV-Hapln1 in CA1 (RM-ANOVA, Virus × Light interaction: *F*_2,19_ = 8.10, *P* < 0.01; no main effect of Virus: *F*_2,19_ = 1.76, *P* = 0.19; main effect of Light: *F*_1,19_ = 16.92, *P* < 0.001). **(R)** Schematic of the contextual fear conditioning protocol. **(S)** P20 mice expressing AAV-Hapln1 in CA1 formed precise contextual fear memories. P20 mice expressing AAV-cfGFP or AAV-ΔHapln1 formed imprecise memories (ANOVA, Virus × Test Context interaction: *F*_2,53_ = 7.83, *P* < 0.01; no main effect of Virus: *F*_2,53_ = 0.027, *P* = 0.97; main effect of Test Context: *F*_1,53_ = 16.62,*P* < 0.001). Data points are individual mice with mean ± s.e.m. Scale bars: white = 50 *μ*m, yellow = 500 *μ*m. * *P* < 0.05; ** *P* < 0.01; *** *P* < 0.001.

Lastly, we tested whether accelerating PNN maturation with AAV-Hapln1 was sufficient to produce adult-like memory phenotypes of sparse engram formation and precise contextual memories (**Fig. 6M-N**). P1 mice were infused with HAPLN1 constructs in the CA1, and we performed optogenetic-mediated allocation and silencing using HSV-NpACY on the same mice on P20-21. Control juveniles (AAV-cfGFP, AAV-ΔHapln1) showed a disruption of allocation to an engram; silencing optogenetically-allocated neurons during the test did not disrupt memory recall. In contrast, juveniles infused with AAV-Hapln1 showed adult-like allocation; silencing optogenetically-allocated neurons during the test disrupted memory recall (**Fig. 6O-Q**). Consistent with this, the engram was sparse in AAV-Hapln1 juveniles and memory was precise. (**Fig. 6R-S, S13P-Q**). Direct infusion of recombinant brain-derived neurotrophic factor (BDNF) protein, a treatment that enhances maturation of neural circuits^32^, into CA1 on P17 also resulted in the precocial maturation of PNNs, PV^+^ interneurons, and memory precision in juvenile mice (**Fig. S16A-I**).

The specific neurobiological mechanisms regulating age-dependent increases in precision of episodic-like memory have long remained elusive^5, 7, 8^. We found that the developing CA1 is actively engaged in early memory formation, despite the immaturity of some memory formation mechanisms. Our findings identify engram sparsity as a key mechanism for memory precision. Specifically, we identified maturation of a competitive neuronal allocation process, supported by developing PV^+^ interneurons, as a prerequisite for encoding precise, episodic-like memories in sparse engrams. These findings support the recent proposal that the hippocampus transitions through distinct stages of functional development, with the stepwise emergence of CA1 phenomena important for memory consolidation^41, 42^. These processes disproportionately affect engram neurons^43, 44^, providing potential mechanisms for post-encoding stabilization of sparse engrams during the fourth postnatal week and onward.

We discovered that maturation of PV^+^ inhibitory circuitry required for the onset of adult-like neuronal allocation and episodic-like memory precision in CA1 are dependent on the extracellular matrix of PNNs. In mammalian and avian brains, the accumulation of PNNs around PV^+^ interneurons in defined cortical regions shifts local excitatory-inhibitory balance, dampens critical period plasticity, and controls the development of corresponding sensory processes or behaviors^31, 35, 37, 39^. Our data suggest that maturation of the hippocampal memory system is regulated by the same cellular and molecular mechanisms as cortical sensory systems. They indicate that memory formation changes the relative levels of perisomatic inhibition of engram neurons through PNN-dependent mechanisms, likely mediated in part by the PNN glycoprotein tenascin-R^45^. Extracellular matrix-dependent maturation of inhibitory neural circuits may be a brain-wide mechanism for not only sensory development, but also cognitive and emotional development^34, 46, 47^.

Why are there multiple stages of functional maturation in the development of the hippocampal memory system? One possibility is that hippocampal development involves progression of the episodic memory system from an ‘incomplete’ (child-like) to ‘complete’ (adult-like) state. An alternative possibility is that the mnemonic functions of children are perfectly adapted for their stage of development^48^. At the cognitive level, the encoding of schemas and other forms of broad or imprecise semantic knowledge during early life may be favored over encoding specific episodes, given that young children have comparatively few life experiences from which to draw and typically are not without an adult caregiver(s) (with fully-fledged episodic memory systems) for the first years of life^49^. The immature hippocampus may have evolved to fulfill this purpose, exploiting the protracted development of inhibition and co-opting the same activity-dependent mechanisms required for structural and functional development of hippocampal circuitry^21–23^ for early memory formation and storage in dense memory engrams.

## Acknowledgments

We thank Antonietta Decristofaro, Daisy Lin, and Mika Yamamoto for their excellent technical assistance. We also thank R. Frischknecht for reagents and the Frankland and Josselyn labs for general comments.

## Funding

Brain Canada (PWF, SAJ)

Canadian Institutes of Health Research (CIHR) grant PJT180530 (PWF) CIHR Postdoctoral Fellowship (AAw)

CIHR Vanier Canada Graduate Scholarship (MLdS)

German Research Foundation (Deutsche Forschungsgemeinschaft) grant SFB1286/A03 (AD)

German Center for Neurodegenerative Diseases (AD)

Hilda and William Courtney Clayton Paediatric Research Fund (SYK)

Hospital for Sick Children Restracomp Fellowship (AAw, LMT)

National Institutes of Health (NIH) grant R01 MH119421 (PWF, SAJ) NIH grant F31 MH120920-01 (AIR)

Natural Sciences and Engineering Research Council of Canada (NSERC) Canada Graduate Scholarship (AIR, LMT)

Ontario Graduate Scholarship (YW)

Ontario Trillium Scholarship (YW)

Vector Institute (LMT)

## Author contributions

Conceptualization: AIR, PWF

Development of HAPLN1 viral constructs: RK, AD Neonatal stereotaxic surgery: AIR

Stereotaxic surgery: AIR, YW, AG, JL

Fear conditioning experiments: AIR, AG, BAY, AJR, LMT, AAb, MA

Spatial foraging task experiments: YW, SYK

Histology and quantification: AIR, YW, AG, BAY, AAb, LCD, CM, JG

Western blot experiments: MLdS, AAw

Statistical analyses: AIR

Funding acquisition: AD, SAJ, PWF

Writing – original draft: AIR, PWF

Writing – review & editing: AIR, AD, SAJ, PWF

## Data and materials availability

The data generated during the current study are available from the corresponding author by reasonable request.

**Supplementary Materials** Materials and Methods Figs. S1 to S16 References (*50*–*67*)

## Materials and Methods

### Mice

Two mouse lines were used. Male and female F1 hybrid (C57BL/6NTac x 129S6/SvEvTac) wild-type (WT) mice were used for all experiments, except where noted. Mice were bred at the Hospital for Sick Children and group-housed on a 12-h light/dark cycle with food and water *ad libitum*. All experiments took place during the light phase. Primiparous and multiparous dams were used for breeding, and litter birthdate was designated postnatal day (P) 0. Litters ranged in size from 5-12 pups. Occasionally, pups were cross-fostered to different lactating dams on P1-P3 to equalize litter sizes and/or increase experiential diversity of offspring. Cross-fostered rodents receive similar maternal care compared with normally-reared offspring^50^. Preweanling mice (Ages ≤ P20) remained in the breeding cage (identical to standard cages) with the dam for the duration of the experiments. Older mice (Ages ≥ P24) were weaned from the dam on P21 and thereafter group-housed with same-sex littermates in standard mouse housing cages (2-5 mice per cage). To rule out weaning as a potential driver of memory development, in a single experiment (see Fig. **S1I-J**), we weaned mice early on P17 (P20 groups) or did not wean mice (P24 groups). Each experimental condition contained mice derived from 2-8 separate litters, with no more than 2 same-sex littermates used for each experimental condition.

PV-Cre knockin driver transgenic mice (B6;129P2-*^Pvalbtm^*^1^(cre)*^Arbr^*/J) which express Cre recombinase in PV^+^ interneurons, without disrupting endogenous PV expression, were originally generated by Silvia Arber, FMI, and obtained from Jackson Laboratory^51^. Homozygous PV-Cre mouse breeders were maintained on a C57BL/6 background and crossed with 129S6/SvEvTac breeders to generate the hybrid PV-Cre mice used in experiments. Housing procedures for this line were identical to those described for WT mice. All procedures were approved by the Hospital for Sick Children Animal Care and Use Committee and conducted in accordance with Canadian Council on Animal Care and National Institutes of Health guidelines.

### Drugs

#### DREADD agonist 21 (C21)

C21 dihydrochloride (Tocris, cat# 6422) was prepared as a stock solution of 10 mg/ml in dH2O and stored at -20 °C. Stock solution was later thawed and diluted 1:10 in PBS. Diluted C21 was administered via i.p. injection 1 h before contextual fear training or recall (1.0 mg/kg) to activate or inhibit DREADD-expressing neurons.

#### Chondroitinase ABC (ChABC)

ChABC from *Proteus vulgaris* (Sigma, cat# C3667) was dissolved in 0.1% bovine serum albumin in PBS at a concentration of 50 U/ml. ChABC solution was stored at -80 °C until use.

#### Penicillinase

Penicillinase from *Bacillus cereus* (Sigma, cat# P0389) was dissolved in 0.1% bovine serum albumin solution (in PBS) at a concentration of 50 U/ml. Penicillinase solution was stored at -80 °C until use.

#### Recombinant Brain-Derived Neurotrophic Factor (BDNF)

Recombinant BDNF (Peprotech, cat# 450-02) was dissolved in PBS at a concentration of 0.36 mg/ml and stored at -80 °C until use.

### Viruses

All viruses were made in-house.

#### Generation of plasmids

To fluorescently label and image extracellular matrix in the noncellular oxidative environment, we subcloned full-length mouse Hapln1 (GeneID: 12950) or ΔHapln1 (mutant lacking the CSPG binding domain) in frame with cysteine-free GFP (cfGFP). The cfGFP plasmid was a gift from Dr. Ikuo Wada^52^. The cDNAs were amplified using the following set of primers: Hapln1 forward primer TAAGCACTCGAGGTGAGCAAGGGCGAGGAGC and reverse primer TAAGCAGGTACCTTACTTGTACAGCTCGTCCATGC; ΔHapln1 forward primer TAAGCACTCGAGGTGAGCAAGGGCGAGGAGC and reverse primer TAAGCAGAGCTCTTACTTGTACAGCTCGTCCATGCCG; cfGFP forward primer

TAAGCAGAGCTCGTGAGCAAGGGCGAGGAGC and reverse primer TAAGCATCTAGATTATTGTACAGCTCGTCCATGCCG and cloned into a custom-designed AAV vector using appropriate restriction enzymes. The complete plasmid sequences can be found in the supplementary materials.

#### HSV

We used replication-defective herpes simplex viruses (HSV) to manipulate sparse subsets of CA1 neurons. HSV is naturally neurotrophic and transfects approximately 30% of principal neurons in CA1 following microinjection (see **Fig. S5**). Transgene expression peaks 3-4 days and dissipates after 10-14 d after HSV microinjection^53^. HSV titers were approximately 1.0 ×10^8^ infectious units/ml. The following HSV constructs were used:

#### HSV-NpACY

We used HSV-NpACY to bidirectionally manipulate the activity of sparse subsets of neurons. HSV-NpACY contains both enhanced channelrhodopsin-2 (ChR2-H134R) fused to enhanced yellow fluorescent protein (eYFP) and halorhodopsin 3.0 (NpHR3.0). These opsins are spectrally compatible, allowing for neuronal excitation by ChR2 with blue light (473 nm) and neuronal silencing by NpHR3.0 with red light (660 nm)^26^. Opsin genes were connected in the viral vector using a 2A self-cleavage linker (p2A) and expression was driven by the endogenous HSV promoter IE4/5.

#### HSV-NpACY-tdTomato

We used HSV-NpACY-tdTomato instead of HSV-NpACY in select experiments in which a different GFP-expressing viral construct was co-expressed in CA1. HSV-NpACY-tdTomato is identical to HSV-NpACY, with the addition of the tdTomato transgene. In this construct, expression of NpACY was driven by the IE4/5 promoter and expression of tdTomato was driven by a downstream CMV promoter.

#### HSV-GFP-hM4Di

We used HSV-GFP-hM4Di to inhibit sparse subsets of CA1 neurons. An hM4Di construct, (a gift from Dr. Bryan Roth, University of North Carolina), was subcloned into an HSV-p1006 vector backbone^54^. In the resulting HSV-GFP-hM4Di construct, expression of GFP was driven by the IE4/5 promoter and expression of hM4Di was driven by the downstream CMV promoter.

#### AAV

We used adeno-associated viruses (AAV) to manipulate neuronal activity or to express our novel Hapln1 constructs. Transgene expression peaks 3-4 weeks after AAV microinjection and is relatively stable in the following weeks. We used a 19-20 d delay allowing roughly equal time for transgene expression following neonatal (P1) or standard surgery protocols. AAVs (DJ serotype) were generated in HEK293T cells with the AAV-DJ Helper Free Packaging System (Cell Biolabs, Inc., cat# VPK-400-DJ) using the manufacturer-suggested protocol. Viral particles were purified using Virabind AAV Purification Kit (Cell Biolabs, Inc., cat# VPK-140). AAV titers were approximately 1.0 ×10^11^ infectious units/ml. The following AAV constructs were used:

#### AAV(DJ)-CaMK2α-iC++-eYFP (AAV-iC++)

We used AAV-iC++ to silence CA1 pyramidal neurons in adult mice. pAAV-CaMK2α-iC++-eYFP (a gift from Dr. Karl Deisseroth) was obtained from Stanford Gene Vector and Virus Core. The chloride channel iC++ enables neuronal silencing with blue light (473 nm) and its expression in excitatory principal neurons was driven by the CaMK2α promoter. We did not use AAV-iC++ for neuronal silencing in juvenile mice because we observed increased locomotion in control (GFP-expressing) juvenile mice receiving constant blue light stimulation during fear recall (see **Fig. S3E-F**).

#### AAV(DJ)-CaMK2α-NpACY (AAV-NpACY)

We used AAV-NpACY to silence CA1 pyramidal neurons in juvenile and adult mice. The transgene is identical to that described for HSV-NpACY, enabling neuronal silencing by NpHR3.0 with red light (660 nm). Expression of NpACY in principal neurons was driven by the CaMK2α promoter.

#### AAV(DJ)-CMV-GFP (AAV-GFP)

We used AAV-GFP as a control virus for AAV-iC++ and AAV-NpACY. pAAV-CMV-GFP was a gift from Dr. Connie Cepko (Addgene plasmid # 67634; http://n2t.net/addgene:67634; RRID:Addgene_67634). Expression of GFP was driven by the CMV promoter.

#### AAV(DJ)-CaMK2α-hM3Dq-mCherry (AAV-hM3Dq)

We used AAV-hM3Dq to activate CA1 pyramidal neurons. pAAV-CaMKIIa-hM3D(Gq)-mCherry was a gift from Dr. Bryan Roth (Addgene plasmid # 50476; http://n2t.net/addgene:50476; RRID:Addgene_50476). Expression of hM3Dq-mCherry in principal neurons was driven by the CaMK2α promoter.

#### AAV(DJ)-hSyn-DIO-hM4Di-mCherry (AAV-DIO-hM4Di)

We used AAV-DIO-hM4Di in PV-Cre mice to inhibit CA1 PV^+^ interneurons. pAAV-hSyn-DIO-hM4D(Gi)-mCherry was a gift from Dr. Bryan Roth (Addgene plasmid # 44362; http://n2t.net/addgene:44362; RRID:Addgene_44362). Expression of hM4Di-mCherry in Cre^+^ interneurons was driven by the neuronal Human synapsin 1 (hSyn) promoter.

#### AAV(DJ)-hSyn-DIO-mCherry (AAV-DIO-mCherry)

We used AAV-DIO-mCherry in PV-Cre mice as a control for AAV-DIO-hM4Di. pAAV-hSyn-DIO-mCherry was a gift from Dr. Bryan Roth (Addgene plasmid # 50459; http://n2t.net/addgene:50459; RRID:Addgene_50459). Expression of mCherry in Cre^+^ interneurons was driven by the neuronal hSyn promoter.

#### AAV(DJ)-hSyn-cfGFP2 (AAV-cfGFP)

We used AAV-cfGFP2 as a control for AAV-Hapln1 and AAV-ΔHapln1. pAAV-Syn-cfGFP2 was a gift from Dr. Ikuo Wada (Fukushima Medical University). Cysteine-free GFP (cfGFP) is a modified GFP that reduces protein oligomerization and restriction of free diffusion in the endoplasmic reticulum. Expression of cfGFP2 was driven by the hSyn promoter.

#### AAV(DJ)-hSyn-Hapln1-cfGFP2 (AAV-Hapln1)

We used AAV-Hapln1 to over-express wild-type mouse HAPLN1 protein with cfGFP2. Expression of Hapln1-cfGFP2 was driven by the hSyn promoter.

#### AAV(DJ)-hSyn-ΔHapln1-cfGFP2 (AAV-ΔHapln1)

We used AAV-ΔHapln1 to over-express mutant (dominant-negative) ΔHAPLN1 protein with cfGFP2. ΔHAPLN1 lacks the wild-type N-terminus IgG domain that binds chondroitin sulfate proteoglycans (CSPGs), while maintaining the C-terminus HA1 and HA2 domains that bind to hyaluronic acid. Expression of Hapln1-cfGFP2 was driven by the hSyn promoter.

### Surgery

Surgeries occurred on P1, P16 and/or P17 (P20 groups), P21 (P24 groups), or P40-onwards (P60 or adult groups). Surgical procedures were similar at all ages, except for the neonatal (P1) timepoint. Mice were pre-treated with atropine sulfate (0.1 mg/kg, i.p.), anesthetized with chloral hydrate (400 mg/kg, i.p.) or isoflurane (3% induction, 1-1.5% maintenance), administered meloxicam (4 mg/kg, s.c.) for analgesia, and placed into stereotaxic frames. The scalp was incised and retracted, and holes were drilled above the dorsal CA1. Unless otherwise specified, viruses or drugs were injected bilaterally via a glass micropipette connected via polyethylene tubing to a microsyringe (Hamilton) at a rate of 0.1 *μ*l/min and remained in place for an additional 10 min to ensure virus diffusion. The following coordinates and virus volumes were used for dorsal CA1. For P16-P17: coordinates AP -1.65 mm, ML ±1.35 mm, DV -1.45 mm from bregma; 0.7 *μ*l HSV; or 0.7 *μ*l BDNF (0.25 ng per side^55^) or vehicle (PBS). For P21: coordinates AP -1.7 mm, ML ± 1.4 mm, DV -1.5 mm from bregma; 0.75 *μ*l HSV. For P40+: coordinates AP -1.8 mm, ML ±1.5 mm, DV -1.5 mm from bregma; 1.0 *μ*l HSV; 0.85 *μ*l AAV; or 1.0 *μ*l ChABC or Penicillinase. In a subset of experiments we microinjected AAVs, ChABC, or Penicillinase into the CA3, dentate gyrus (DG), prelimbic cortex (PrL), or retrosplenial cortex (RSC) of adult mice using identical procedures. For CA3: coordinates AP -2.2 mm, ML ± 2.7 mm, DV -2.5 mm from bregma; 0.85 *μ*l AAV. For DG: coordinates AP -2.2 mm, ML ±1.65 mm, DV -2.2 mm from bregma; 0.85 *μ*l AAV. For PrL: coordinates AP +1.7 mm, ML ±0.35 mm, DV -1.8 mm from bregma; 0.65 *μ*l AAV at a rate of 0.05 *μ*l/min. For RSC: coordinates AP -1.5 mm, ML ±0.35 mm, DV -1.15 mm from bregma; 1.0 *μ*l ChABC or Penicillinase at a rate of 0.05 *μ*l/min. For western blot experiments, we targeted all subfields of the dorsal hippocampus using coordinates AP -2.0, ML ±2.0, DV -2.2 and -1.5 mm. For each DV site, 1.0 *μ*l of AAV, ChABC, or Penicillinase was injected. Following both microinjections, the scalp was sutured and polysporin was applied to the wound. Mice were administered 0.9% saline (0.5-1.0 ml, s.c.) and placed in a clean cage on a heating pad to recover. Once recovered, the dam was moved to the clean cage (for P16-17 mice), or mice were weaned immediately (for P21 mice).

Neonatal virus injection procedures were adapted from previous studies^21, 56^. P1 mice were anesthetized through hypothermia and mounted to a chilled metal plate using laboratory tape. The scalp was incised and retracted, and the glass micropipette connected to a nanoliter injector (Nanoject III, Drummond Scientific) was used to pierce the skull above dorsal CA1 (approximate coordinates AP -0.5 mm, ML ±0.8 mm from bregma). The pipette was lowered to DV -1.1 from the skull surface, and 0.12-0.15 *μ*l AAV was injected over 2 min and remained in place for an additional 2 min. The scalp was sealed with Vetbond Tissue Adhesive (3M) and covered with polysporin. For some experiments, paws were tattooed using non-toxic black ink for later mouse identification^57^. The entire procedure was performed within 10-12 min. Pups were placed on a heating pad and continuously monitored until mobile. Pups remained on the heating pad until all surgeries were completed, at which time the litter was returned to the dam.

For optogenetics experiments, optic fibers were implanted above the dorsal CA1 on P17 (P20 groups), P21 (P24 groups), or P53-57 (P60 groups). Implants were constructed in-house by attaching a piece of polished 200 *μ*m diameter optic fiber (0.37 numerical aperture) to a 1.25-mm diameter zirconia ferrule with epoxy resin. Optical fiber implantation was performed immediately following virus injection for HSV experiments or in a second procedure for AAV experiments. Fiber tips were lowered to DV -1.2 mm from bregma above the dorsal CA1 and secured to the skull using screws and black dental cement. Post-surgery procedures and care were the same as described above.

For medial forebrain bundle (MFB) stimulation experiments, unilateral MFB stimulation was conducted as previously described^58^. MFB implants were performed on P16 (P20 groups), P21 (P24 groups), or P53 (P60 groups). Concentric bipolar electrodes were lowered into the right MFB using the following stereotaxic coordinates. For P16 and P21: coordinates AP -1.1 mm, ML 1.0 mm, DV -5.1 mm from bregma. For P53: coordinates AP -1.2 mm, ML 1.0 mm, DV -5.2 mm from bregma. Implants were attached to the skull using screws and black dental cement. Correct placement of the electrodes in the MFB was verified during surgery by brief electrical pulses and by post-hoc identification of electrode tracks in brain slices. Post-surgery procedures and care were the same as described above.

For all experiments involving virus microinjections, only mice showing strong bilateral expression (i.e., expression limited to the target brain region and observable in a minimum of 3 brain sections) were included in the final data set. For experiments in which we infused a combination of HSV and/or AAV constructs, only mice correctly expressing both transgenes in the same region were included in the final data set. Additionally, for optogenetics experiments, only mice with optic fibers placed correctly above the opsin-expressing region of interest were included in the final data set.

### Behavior

#### Fear conditioning

Contextual fear conditioning occurred in test chambers (31 × 24 × 21 cm) with shock-grid floors (Med Associates). Unless otherwise stated, mice were trained in a single 5-min session with three foot shocks. During training, mice were placed in the chambers for 2 min, after which three foot shocks (0.5 mA, 2 s duration, 1 min apart) were delivered. Mice were removed from the chambers 1 min after the last shock. The next day, mice were placed in one of four different test chamber configurations for 5 min or 6 min (for optogenetics experiments). In most experiments, testing occurred in the training chamber (Context A) or a similar but novel chamber (Context B). Context B was approximately the same size as Context A, with white plastic floor and triangular white plastic walls. Two novel, dissimilar contexts were used for specific experiments. Context C was a rectangular chamber (15 × 45 × 25 cm) with an open top, made of white plexiglass, and was located in a separate room. Context D was a test chamber with white plastic floor and a semi-circular white plastic wall, with speakers for auditory tone presentation. Context D chambers were located in another separate testing room and placed inside of sound-attenuating boxes. Mouse behavior was recorded with overhead cameras and FreezeFrame v.3.32 software (Actimetrics). For contextual fear memory tests, the amount of time mice spent freezing was scored during the entire test session automatically using FreezeFrame software or manually (for optogenetics experiments). Freezing was defined as the cessation of movement, except for breathing^59^. Specific details for experiments deviating from the standard protocol are described below.

To test whether juvenile mice’s potentially different learning rate or training intensity accounts for their memory imprecision, we modified the training protocol for P20 and P60 mice to account for differences in freezing after training (see **Fig. S1A-G**). After 2 min in the chamber, P20 mice received one foot shock (0.5 mA, 2 s duration) or P60 mice received 5 foot shocks (0.5 mA, 2 s duration, 1 min apart), and were removed after 1 min. Testing occurred as described above. To test whether juvenile mice’s lack of familiarity with the test chambers accounts for their memory imprecision, we pre-exposed mice to chambers before fear conditioning (see Fig. S1K-O). Context pre-exposure is an experiential manipulation known to increase memory precision in adult rats and mice^60, 61^. Mice were pre-exposed to Contexts A and B (order counterbalanced) on P19 (P20 groups) or P23 (P24 groups) for 5 min each with 5-6 h between exposures (Pre groups). Control mice were not pre-exposed to either context before training (No-Pre groups). Training and testing began the following day as described above.

To test whether juvenile mice formed imprecise short-term memories, we reduced the time interval between training and testing (see **Fig. S1R-S**). P20 and P60 mice were tested 1 h after training.

To test whether memory imprecision in juvenile mice extended to all novel environments, we tested mice in a novel, dissimilar chamber (see **Fig. S1T-U**). Following training, P20, P24, and **Fig. S13C-D** P60 mice were tested in Context C.

To test whether the effects of CA1 PNN disruption were specific to contextual and spatial memories, we trained adult mice expressing ΔHapln1 or cfGFP in auditory fear conditioning (see) as previously described^26^. During training, mice were placed in the conditioning chamber for 2 min before the presentation of a 30 s auditory tone (2.8 kHz tone, 85 dB; Tone A) that co-terminated with a foot shock (0.5 mA, 2 s duration). Mice remained in the chamber for an additional 30 s after the shock. For the test, mice were placed into Context D, and after 2 min, presented with Tone A or a novel auditory stimulus (7.5 kHz pips, 5 ms rise, 75 dB; Tone B) for 1 min. Freezing behavior to Tone A and Tone B were scored automatically.

#### Optogenetic stimulation (contextual fear conditioning)

Mice used in the optogenetics experiments were habituated to the optic patch cables for 5 min on the day before training. For optogenetic-mediated allocation experiments, the allocation procedure was performed immediately before training in the contextual fear conditioning task. Mice were attached to the optic patch cables and placed into a clean cage. NpACY^+^ neurons in these mice were briefly excited with blue light (473 nm, 1 mW, 4 Hz, 15-ms pulses) for 30 s. This stimulation frequency was chosen based on previous reports^18, 62^. Non-allocated, Control mice did not receive light stimulation. Mice were detached from the optic patch cables and trained immediately as described above. We verified that optogenetic-mediated allocation before training results in increased localization of training-induced c-Fos in NpACY^+^ neurons (see **Fig. S5**). For optogenetic stimulation during testing, mice were attached to the patch cables and placed in the test chamber (Context A or Context B) for a 6 min period. For the first 3 min of the test session, no light was applied. For the latter 3 min of the test, continuous blue (473 nm, 7-10 mW) or red (660 nm, 7-10 mW) light stimulation was delivered to silence the iC++^+^ or NpACY^+^ neurons in CA1. Freezing behavior during these memory tests was scored manually by an experimenter blinded to the experimental conditions (except Age, which could not be blinded).

#### Exploratory habituation

To determine whether juvenile mice could perceptually discriminate similar spatial environments, we performed a non-associative exploratory habituation task (see **Fig. S1P-Q**). P20 and P60 mice were placed in a rectangular shuttle box (15 × 45 × 25 cm) with white and black-and-white striped compartments and allowed to freely explore for 5 min (Exposure 1). 24 h later, mice were placed into the same shuttle box (Context A) or a different shuttle box with light and dark compartments (Context B) for 5 min (Exposure 2). Mouse behavior was recorded using overhead cameras and the distance traveled during the 5-min sessions was scored automatically using Limelight software (Actimetrics). Because distance traveled was overall lower for P20 mice compared to P60 mice (data not shown), we quantified exploratory habituation to the environments by computing a Habituation Index (Exposure 2 Distance / Exposure 1 Distance) where a score of 1.0 indicates no change in exploration and scores less than or more than 1.0 indicate exploratory habituation or sensitization, respectively.

#### Inhibitory Avoidance (IA)

Inhibitory avoidance was performed as previously described^55^. The IA apparatus was a rectangular shuttle box (25.5 × 16.5 × 17.7 cm) designed similarly to the shuttle boxes used for exploratory habituation experiments. The IA shuttle box consisted of two compartments (safe and shock compartments) separated by a sliding door. The safe compartment was white and illuminated and the shock compartment was black and covered to prevent light entry. On the training day, mice were placed in the safe compartment with their heads facing away from the closed door, and after 10 s, the door was opened. The sliding door was shut 1 s after mice entered the shock compartment, and a single 1 mA (2 s duration) foot shock was delivered. Mice remained in the shock compartment for 10 s following the foot shock and then were returned to their home cage. The next day, mice were tested using the training IA shuttle box (Context A) or a modified shuttle box (Context B). The walls of the Context B shuttle box were curved and Context B similarly contained light- and dark-colored compartments separated by a sliding door. Mice were placed in one of the test apparatuses in the illuminated compartment, and after 10 s the sliding door was opened. The latency (in s) of the mice to enter the dark compartment from the time of door opening was examined for the training and testing sessions.

#### Spatial Foraging Task.

The spatial foraging task was designed similarly to the hippocampus-dependent Morris water maze task^63^. In contrast to the Morris water maze task, performance in the spatial foraging task is appetitively-motivated and allows for easier manipulation of the testing environments. Mice were handled and habituated to the stimulation patch cord for 15 min/day for 3 (P20 and P24 groups) or 7 (P60 groups) days before training. Training occurred in a white, square arena (100 × 100 × 40 cm for P60 groups or 42 × 42 × 30 cm for P20 and P24 groups) located in a dimly lit room. The arena was surrounded by white curtains painted with four distinct visual cues, located 1 m away from the perimeter of the arena. The arena was divided into quadrants, and a circular reward zone (11 cm in diameter for P60 groups and 6 cm in diameter for P20 and P24 groups) was located in the center of one of the four quadrants during training. On the training day, mice received 20 training trials that were 3 min in duration each. Electrical stimulation of the MFB was designed to approximate high-frequency brain stimulation (139 Hz, 90-μs pulses). During the first trial, mice were connected to the stimulating patch cord, placed into the reward zone, and MFB stimulation was delivered. This was done to expose mice to MFB stimulation within the arena’s reward zone. On subsequent trials, mice were placed in one of four pseudo-randomly chosen starting locations and allowed to explore freely. MFB stimulation was delivered upon entry into the reward zone and was terminated once mice left the reward zone. If mice failed to find the reward zone within 60 s from the trial start, they were gently guided by the experimenter to the location. Twenty-four-h later, a 60-s probe trial was conducted in the training arena (Context A) or a novel circular arena (Context B) placed in the same position as the training arena (i.e., with the same distal visual cues). Mice were placed in the quadrant furthest from the reward zone during training and allowed to freely explore the testing arena, with no delivery of MFB stimulation. In one experiment, we removed the visual cues from the curtain surrounding the testing arena to demonstrate that performance in the spatial foraging task (like the Morris water maze) is dependent on the use of distal spatial cues (see **Fig. S2O-Q**).

Behavioral data from the spatial foraging task were acquired and analyzed using custom software. For training, latency to reach the reward zone (in seconds, maximum of 60 s) was recorded. For the probe trial, we quantified spatial memory precision by measuring time in quadrants, proximity to the reward zone, and distribution of angular positions. For the primary quadrant analysis, the amount of time mice spent searching in the arena quadrant that previously held the reward zone in the training Context A or the equivalent quadrant in the novel Context B (chance performance = 25%). For the proximity analysis, the linear distance between the mouse body and center of the reward zone (in cm) was calculated for every time bin. For the angular position analysis, the angle (in radians) between a vector separating the North-East and South-East quadrants of the arena (0 rads) and a vector directed towards the mouse body was calculated for every time bin. Proximity and angular position data were averaged across time to obtain a single data point per mouse (mean proximity and mean angular position).

#### Open field and Elevated-plus maze

To control for the possibility that PNN manipulations alter locomotion or anxiety-like behaviors, we examined mouse behavior in an open field and on an elevated plus maze. The open field was a square arena (45 × 45 × 20 cm) located in a dimly lit room. Mice were placed in the center of the arena and allowed to explore for 10 min. The locomotion of the mice was tracked using an overhead camera. The total distance traveled (normalized to the mean distance of the control group) and amount of time spent in the center of the arena were obtained with Limelight software (Actimetrics). The next day, mice were placed in the center of the elevated plus maze and behavior was monitored for 5 min. The total distance traveled (normalized to the mean distance of the control group) and amount of time spent in the open arms of the maze were obtained using an overhead camera and Ethovision software (Noldus). For both the open field and elevated-plus maze, a reduction in distance traveled or time spent in the center or open-arm regions are considered to reflect an anxiety-like phenotype.

### Histology

#### Perfusion and tissue preparation

At the appropriate age, following the appropriate delay (for viral or drug manipulations), or following behavior experiments, mice were transcardially perfused with chilled 1× PBS followed by 4% paraformaldehyde (PFA), fixed in PFA overnight at 4 °C, and transferred to 30% sucrose solution for at least 48 h. PBS and PFA volumes were adjusted for different mouse ages. When appropriate, perfusions were timed to occur 90 min or 24 h after contextual fear conditioning training or testing. Brains were sectioned coronally using a cryostat (Leica CM1850), and a ¼ sampling fraction was used to obtain 4 sets of 50-μm sections. Sections for immunohistochemistry were stored in 0.1% NaN3 solution until staining.

#### Immunohistochemistry

Immunofluorescence staining was conducted as previously described^26^. For experiments involving quantification of the number of cells positive for immunofluorescence or immunofluorescence signal intensity, all staining was performed at once using the same antibody solutions. Free-floating sections were blocked with PBS containing 4% normal goat serum and 0.5% Triton-X for 1 h at room temperature. Afterwards, sections were incubated with primary antibodies in fresh blocking solution: rabbit anti-c-Fos (1:1000, Synaptic Systems, cat# 226003), rabbit anti-Arc (1:1000, Synaptic Systems, cat# 156003), chicken anti-GFP (1:1000, Aves, cat# GFP-1010), mouse anti-RFP (1:1000, Rockland Immunochemicals, cat# 200-301-379), rabbit anti-RFP (1:1000, Rockland Immunochemicals, cat# 600-401-379), mouse anti-PV (1:1000, Sigma, cat# 1572), rabbit anti-PV (1:1000, Swant, cat# PV27), guinea-pig anti-PV (1:1000, Swant, cat# GP72), mouse anti-Syt2 (1:1000, ZIRC, cat# znp-1), mouse anti-CaMK2ɑ (1:200, Sigma, cat #C265), biotinylated WFA (1:1000, Sigma, cat# L1516), rabbit anti-BCAN (1:1000, a gift from Dr. Renato Frischknecht), and mouse anti-HAPLN1 (1:100, R&D Systems, cat# MAB2608) for 24 h or 72 h (for c-Fos experiments) at 4 °C. Sections were washed three times for 15 min each with PBS containing 0.1% Tween-20 (PBS-T), then incubated with PBS-T containing secondary antibodies: goat anti-chicken Alexa Fluor 488 (1:500, Invitrogen, cat# A-11039, goat anti-mouse Alexa Fluor 488 (1:500, Invitrogen, cat# A-11001), goat anti-mouse Alexa Fluor 568 (1:500, Invitrogen, cat# A-11004), goat anti-mouse Alexa Fluor 647 (1:500, Invitrogen, cat# A-21235), goat anti-rabbit Alexa Fluor 488 (1:500, Invitrogen, cat# A-11008), goat anti-rabbit Alexa Fluor 568 (1:500, Invitrogen, cat# A-11011), goat anti-mouse Alexa Fluor 647 (1:500, Invitrogen, cat# A-21244), goat anti-mouse Alexa Fluor 488 (1:500, Invitrogen, cat# A-11001), goat anti-guinea pig Alexa Fluor 647 (1:500, Invitrogen, cat# A-21450), streptavidin Alexa Fluor 488 (1:500, Invitrogen, cat# S-11223), streptavidin Alexa Fluor 568 (1:500, Invitrogen, cat# S-11226), streptavidin Alexa Fluor 647 (1:500, Invitrogen, cat# S-32357) for 2 h at room temperature. Sections were washed with PBS, counterstained with DAPI, mounted on gel-coated slides, and coverslipped with Permafluor mounting medium (ThermoFisher Scientific, cat# TA-030-FM).

#### Imaging

Images were obtained using a confocal laser scanning microscope (LSM 710; Zeiss). For visualization of virus expression, images were acquired with a 20× objective. For image quantification, z-stacks were acquired using a 40× objective (N.A. = 1.3; 15-40 slices with a 1 *μ*m step size), except for one set of experiments in which images acquired with the 20× objective were used for quantification (see **Fig. S9**). For all experiments involving quantification of the number of cells positive for immunofluorescence or immunofluorescence signal intensity, all images were acquired using identical imaging parameters (laser power, photomultiplier gain, pinhole, and detection filter settings) in a minimal number of imaging sessions (when possible, in one session). For each experiment, imaging parameters were set using a sample section from a control mouse. In experiments where the dependent variable was mouse age, P60 was considered the ‘control’ group.

#### Quantification

For cell counting experiments in the CA1, CA3, or DG, every fourth section was assessed for the marker(s) of interest. For each mouse, cells were counted manually in Fiji (National Institutes of Health) using 5 images acquired from 3-5 sections and averaged. To estimate the number of DAPI^+^ cells in the CA1 pyramidal layer, DAPI^+^ cells were counted in a small volume (approximately 12-20 ×10^3^ *μ*m^3^) to obtain the DAPI^+^ density (mm^-3^) for each Age group (see **Fig. S4**). For all experiments, the volume of the pyramidal layer within each image was measured and multiplied by the DAPI^+^ density value for the appropriate age group to obtain the estimated DAPI^+^ cell number. Quantification of c-Fos^+^ cells was limited to the pyramidal layer, whereas all other markers were quantified in all visible layers. The proportion of pyramidal layer cells expressing c-Fos after contextual fear conditioning training, test, or in the home cage are reported as *P*(c-Fos^+^|DAPI^+^). The proportion of NpACY^+^ cells expressing c-Fos after contextual fear conditioning training in Allocation and No Allocation groups are reported as *P*(c-Fos^+^|NpACY^+^). The number of PV^+^ interneurons and WFA^+^, BCAN^+^, or HAPLN1^+^ PNNs are reported per volume (mm^-3^). Colocalization of the same PNN markers with PV^+^ interneurons are reported as *P*(marker^+^|PV^+^).

To examine how PV^+^ interneuron processes mature with age or are affected by PNN manipulations, we quantified the density of PV^+^ neurites or Syt2^+^ synaptic puncta labeling within the CA1 pyramidal layer^64, 65^. PV^+^ or Syt2^+^ immunofluorescence was binarized in Fiji using a gray value threshold that was manually determined using an image from a mouse in the control or P60 group. The same threshold was applied to all images belonging to the same experiment. We measured the percentage of pyramidal layer area covered by PV^+^ or Syt2^+^ labeling in 15 ROIs per mouse (3 per image, approximately 400 *μ*m^2^ each) and averaged the data to obtain a single data point. For each image (z-stack), ROIs were drawn in z-slice with the greatest immunofluorescence and excluded any areas containing PV^+^ somas.

To examine the structural plasticity of PV^+^ neurites after contextual fear conditioning training, we quantified the density of PV^+^ neurite in the perisomatic region of CA1 pyramidal layer cells, similar to previous studies^26, 66, 67^. PV^+^ immunofluorescence was binarized in Fiji as described above. To quantify the extent of perisomatic PV innervation surrounding c-Fos^+^ and an equal number of c-Fos^-^ cells in the pyramidal layer (50-100 cells per mouse), the DAPI^+^ nucleus of cells were outlined and the percentage of area covered by PV^+^ labeling within a 3-μm band was measured. c-Fos^+^ and c-Fos^-^ cells were approximately matched by selecting c-Fos^-^ cells in close proximity to (within 2 cells away) and adjacent to (and not above or below) a corresponding c-Fos^+^ cell. Perisomatic PV^+^ labeling surrounding c-Fos^+^ and c-Fos^-^ cells were averaged independently for each mouse and used to compute the Perisomatic PV^+^ Selectivity (mean c-Fos^-^ labeling / mean c-Fos^+^ labeling). As a control experiment, we performed the same analysis using P20 and P24 mice that were not trained, except using 20-80 cells per mouse, as the number of c-Fos^+^ cells in P24 mice taken from the home cage was very low.

### Western Blot

Three weeks following AAV microinjections or 24 h following ChABC or Penicillinase microinjections, mice were rapidly decapitated, and their hippocampi were dissected and flash frozen in liquid nitrogen. Dorsal hippocampi were sonicated in homogenization buffer (50 mM Tris-HCl pH 7.5, 0.25 M sucrose, 25 mM KCl, 5 mM MgCl2) supplemented with protease inhibitor cocktail (BioShop, cat# PIC002). Homogenized samples were centrifuged at 14,000 rpm for 15 min at 4°C and the protein concentration of the supernatant was determined by Pierce BCA Protein Assay (ThermoFisher Scientific, cat# 23227). Samples were diluted and supplemented with SDS sample buffer (50 mM Tris-HCl pH 6.8, 2% SDS, 1% β-mercaptoethanol, 5% glycerol, bromophenol blue). Protein samples (25 µg) were separated by electrophoresis on 4-15% mini-PROTEAN TGX precast gels (Bio-Rad, cat# 456-1083) and transferred to PVDF membranes (Bio-Rad, cat# 162-0177). Membranes were blocked with 5% skim milk in Tris-buffered saline with 0.1% Tween-20 (TBS-T) for 1 h at room temperature. Membranes were incubated in primary antibodies diluted in 5% skim milk in TBST: mouse anti-brevican (1:1000, BioLegend, cat# 820101), sheep anti-neurocan (1:1000, R&D Systems, cat# AF5800), goat anti-Hapln1 (1:1000, R&D Systems, cat# AF2608), rabbit anti-β-actin (1:1000, Cell Signaling Technology, cat# 4967) overnight at 4 °C or for 1 h at room temperature. Membranes were incubated with horseradish peroxidase (HRP)-conjugated secondary antibodies diluted in 5% skim milk in TBST: horse anti-mouse IgG (1:20,000, Cell Signaling Technology, cat# 7076), donkey anti-sheep IgG (1:20,000, Invitrogen, cat# A16041), donkey anti-goat IgG (1:20,000, Invitrogen, cat# A15999), and goat anti-rabbit IgG (1:20,000, Sigma, cat# A0545) for 1 h at room temperature. Protein bands were visualized with Amersham ECL Prime Western Blotting Detection Reagent (Cytiva, cat# RPN2236) and chemiluminescence was imaged with a ChemiDoc XRS+ System (Bio-Rad). Band intensities were quantified using Image Lab 6.1 software (Bio-Rad). The intensities of the proteins of interest were normalized to the intensity of the loading control, actin, and normalized to the mean intensity of the control condition in each experiment (AAV-cfGFP or Penicillinase).

### Statistical Analyses

No statistical tests were used to predetermine sample sizes, but our sample sizes were similar to those reported in previous publications^26, 27, 53^. Data were analyzed using one-way or factorial analysis of variance (ANOVA) with repeated measures when appropriate. When appropriate, ANOVAs were followed by Newman-Keuls post-hoc comparisons. In one instance, we used planned *t*-tests based on strong *a priori* predictions (see Fig. 5L). For the perisomatic PV^+^ selectivity analyses in trained (see Fig. 3F) and naive (see **Fig. S7C**) mice, one-sample *t*-tests were used to compare data to a hypothetical mean of 1 (i.e., no change in selectivity). In some experiments, we pooled control mice into a single control group (see **Fig. S8M-N** and **S13J-K**). Sub-group means were not statistically different from one another (data not shown). For the analyses of potential sex differences in memory precision or PNN development (see **Fig. S1H**, **S10A**), data were pooled from different experiments (Fig. 1C**, S1M** (No Pre Group), **S1O** (No Pre Group) and Fig. 4C**, S10C**) to attain sufficiently large sample sizes, and re-analyzed. Subjects pooled into the same groups for the analysis of potential sex differences were treated identically across the different experiments. Statistical significance was set at *P* < 0.05, and Bonferroni correction was applied when appropriate. Statistical analyses were performed using Statistica software (Dell Inc. 2016, version 13) and Graphpad Prism (version 8.0.1).

---

**Figure S1.**
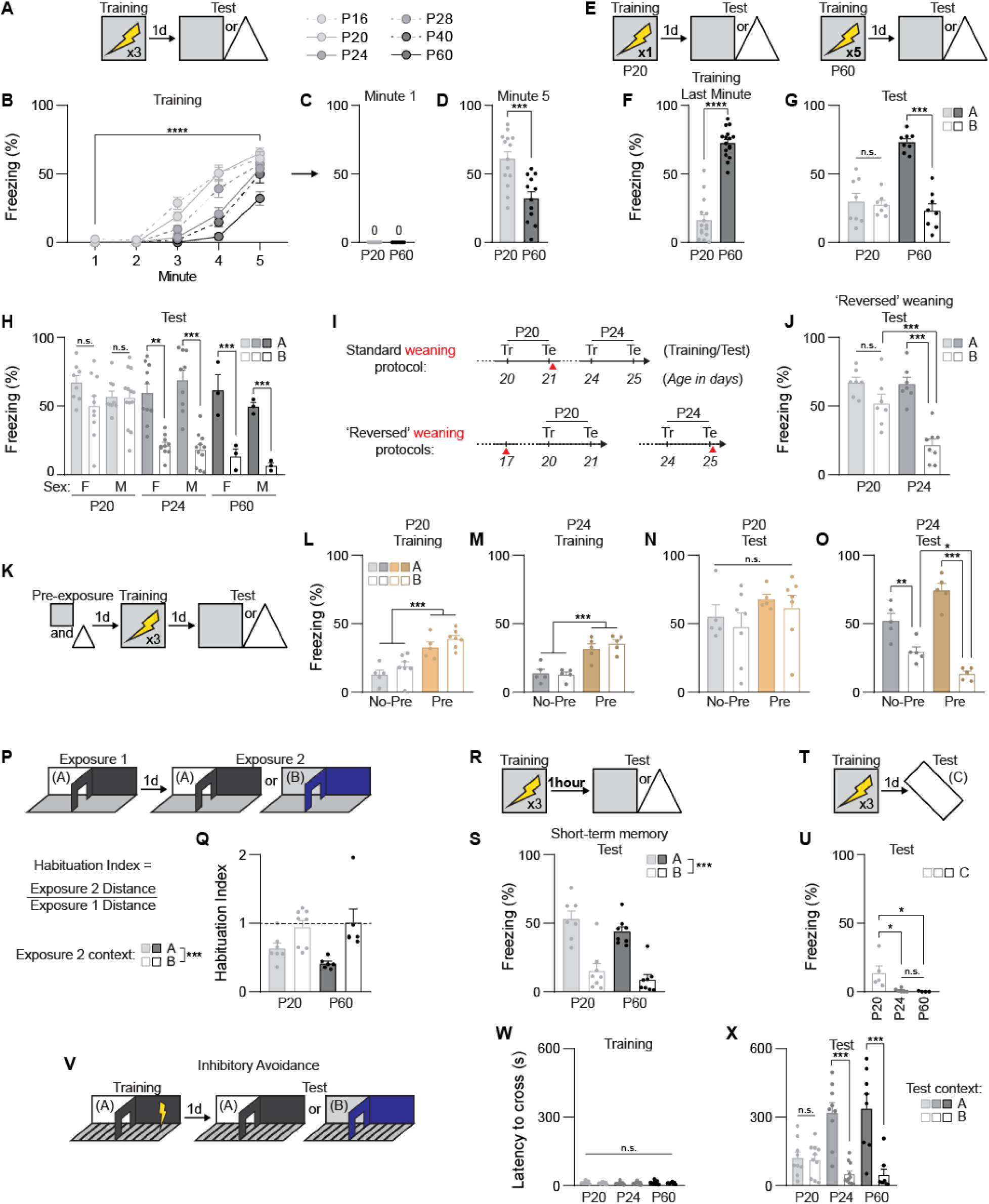
Behavioral characterization of contextual fear memory precision development (Related to Figure 1). **(A to D)** Training data related to Figure 1C. Schematic of the contextual fear conditioning protocol (A). All age groups showed increased freezing across the 5-min training session (B, ANOVA, Age × Minute interaction: *F*_20,304_ = 6.81, *P* < 0.000001; main effect of Age: *F*_5,76_ = 14.66, *P* < 0.000001; main effect of Minute: *F*_4,304_ = 294.63, *P* < 0.000001). P20 and P60 mice exhibited no freezing behavior during the first minute of the training session (C), but P20 mice froze more than P60 mice during the last minute of the training session (D, unpaired *t*-test: *t*_24_ = 4.01, *P* < 0.001). **(E to G)** The contextual fear conditioning protocols were modified to account for potential age-related differences in learning (E). Decreasing and increasing the number of shocks delivered to P20 and P60 mice, respectively, resulted in greater freezing in P60 mice during the last minute of the training session (F, unpaired *t*-test: *t*_29_ = 12.09, *P* < 0.0001), but did not alter memory precision at either age (G, ANOVA, Age × Test Context interaction: *F*_1,27_ = 29.46, *P* < 0.0001; main effect of Age: *F*_1,27_ = 19.78, *P* < 0.001; main effect of Test Context: *F*_1,27_ = 35.25, *P* < 0.00001). **(H)** The development of memory precision by P24 was not sex-dependent (ANOVA, no Age × Sex × Test Context interaction: *F*_2,80_ = 1.77, *P* = 0.17; no Age × Sex interaction: *F*_2,80_ = 0.69, *P* = 0.50; Age × Test Context interaction: *F*_2,80_ = 12.45, *P* < 0.0001; no Sex × Test Context interaction: *F*_1,80_ = 0.12, *P* = 0.72; main effect of Age: *F*_2,80_ = 13.50, *P* < 0.00001; no main effect of Sex: *F*_1,80_ = 0.46, *P* = 0.49; main effect of Test Context: *F*_1,80_ = 64.12, *P* < 0.000001). **(I to J)** Weaning status in P20 and P24 groups (I), did not alter the development of memory precision (J, ANOVA, Age × Test Context interaction: *F*_1,25_ = 8.48, *P* < 0.01; main effect of Age: *F*_1,25_ = 9.85, *P* < 0.01; main effect of Test Context: *F*_1,25_ = 35.25, *P* < 0.00001). **(K to O)** Pre-exposure to the testing contexts (K) improved encoding of fear memories in P20 (L, ANOVA, no Pre-exposure × Test Context interaction: *F*_1,20_ = 0.0010, *P* = 0.97; main effect of Pre-exposure: *F*_1,20_ = 37.70, *P* < 0.00001; no main effect of Test Context: *F*_1,20_ = 3.41, *P* = 0.07) and P24 (M, ANOVA, no Pre-exposure × Test Context interaction: *F*_1,16_ = 0.66, *P* = 0.42; main effect of Pre-exposure: *F*_1,16_ = 48.90, *P* < 0.00001; no main effect of Test Context: *F*_1,16_ = 0.24, *P* = 0.62) mice. Context pre-exposure did not ameliorate memory precision in P20 mice (N, ANOVA, no Pre-exposure × Test Context interaction: *F*_1,20_ = 0.0034, *P* = 0.95; no main effect of Pre-exposure: *F*_1,20_ = 2.11, *P* = 0.16; no main effect of Test Context: *F*_1,20_ = 0.58, *P* = 0.45) but improved memory precision in P24 mice (O, ANOVA, Pre-exposure × Test Context interaction: *F*_1,16_ = 19.90, *P* < 0.001; no main effect of Pre-exposure: *F*_1,16_ = 0.50, *P* = 0.48; main effect of Test Context: *F*_1,16_ = 94.31, *P* < 0.000001). **(P to Q)** P20 and P60 mice successively exposed to the same or different similar environments (P) showed equal levels of exploratory habituation and dishabituation (Q, ANOVA, no Age × Exposure 2 Context interaction: *F*_1,23_ = 1.69, *P* = 0.20; no main effect of Age: *F*_1,23_ = 0.45, *P* = 0.50; main effect of Exposure 2 Context: *F*_1,23_ = 16.64, *P* < 0.001). **(R to S)** Shortening the retention interval between training and testing to 1 hour (R) revealed precise short-term contextual fear memories in P20 mice (S, ANOVA, no Age × Test Context interaction: *F*_1,27_ = 0.10, *P* = 0.75; no main effect of Age: *F*_1,27_ = 2.83, *P* = 0.10; main effect of Test Context: *F*_1,27_ = 62.13, *P* < 0.000001). **(T to U)** P20 mice tested in a environment dissimilar to the training context (T) showed little freezing behavior, but froze more than P24 and P60 mice (U, ANOVA, main effect of Age: *F*_2,12_ = 6.17, *P* < 0.05). **(V to X)** In an inhibitory avoidance task (V), all mice had similar crossing latencies during training (W, ANOVA, no Age × Test Context interaction: *F*_2, 49_ = 0.62, *P* = 0.53; no main effect of Age: *F*_2,49_ = 0.19, *P* = 0.82; no main effect of Test Context: *F*_1,49_ = 0.69, *P* = 0.40). During testing, P20 mice had imprecise inhibitory avoidance memories compared to P24 and P60 mice, which showed lower crossing latencies in the novel shuttle box (X, ANOVA, Age × Test Context interaction: *F*_2, 49_ = 10.78, *P* < 0.001; no main effect of Age: *F*_2,49_ = 2.95, *P* = 0.06; main effect of Test Context: *F*_1,49_ = 45.77, *P* < 0.000001). Data points are individual mice with mean ± s.e.m. * *P* < 0.05; ** *P* < 0.01; *** *P* < 0.001, **** *P* < 0.0001.

**Figure S2.**
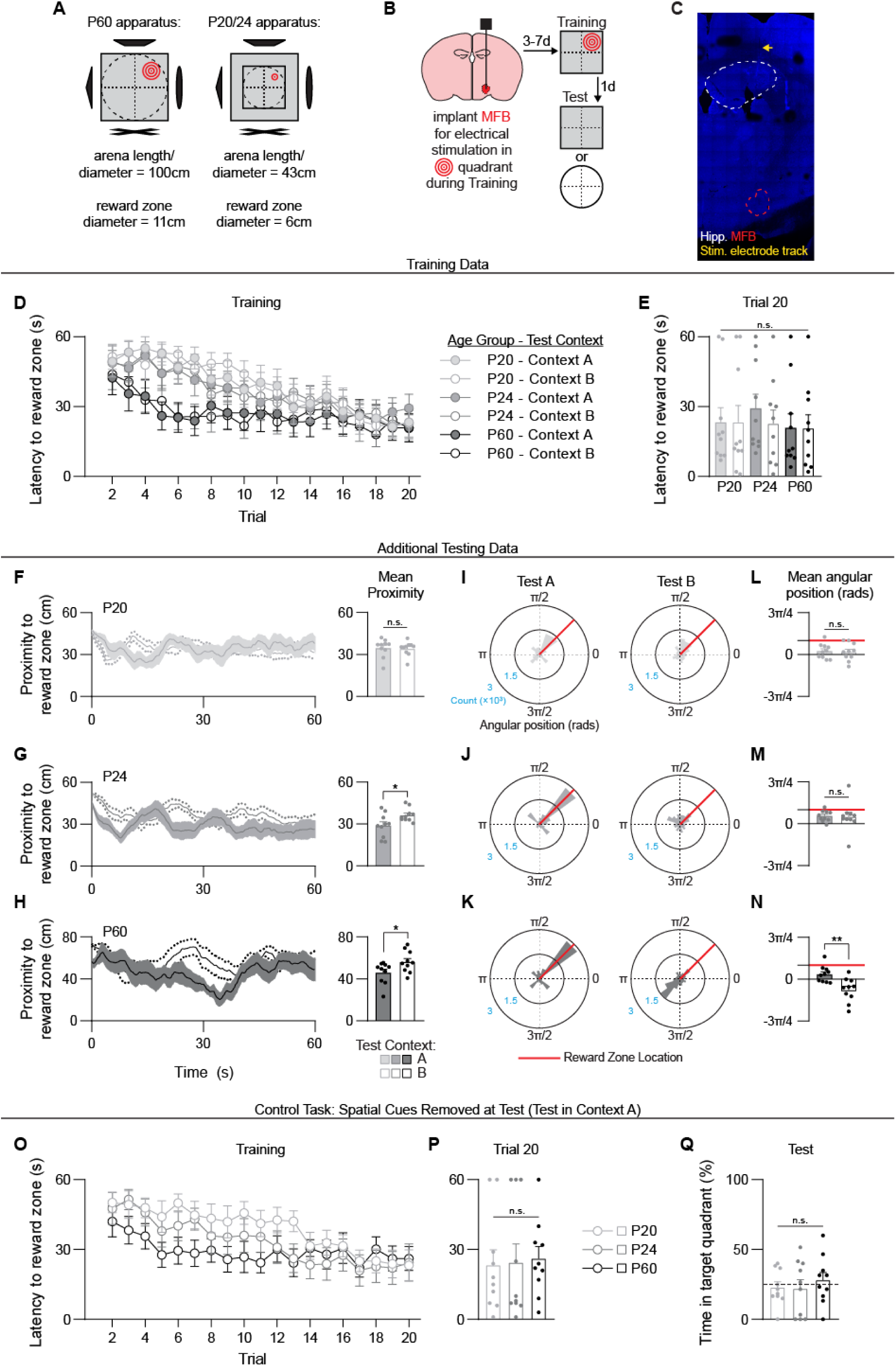
Spatial foraging task training data, additional test performance measures, and control task (Related to Figure 1). **(A)** Schematic of testing apparatuses used for the spatial foraging. **(B)** Mice were implanted with stimulating electrodes targeting the medial forebrain bundle (MFB) days before training and testing. **(C)** Representative image showing electrode tracks above the MFB in the brain of a P60 mouse. **(D to E)** Latencies to locate the reward zone decreased across training trials in all groups (D) and did not differ across groups by the last training trial (E, ANOVA, no Age × Test Context interaction: *F*_2,54_ = 0.18, *P* = 0.83; no main effect of Age: *F*_2,54_ = 0.33, *P* = 0.71; no main effect of Test Context: *F*_1,54_ = 0.65, *P* = 0.65). **(F to H)** During the test, the average proximity to the previously rewarded zone in Context A or the equivalent area in Context B did not differ in P20 mice (F, unpaired *t*-test: *t*_18_ = 0.17, *P* = 0.86), but was lower in Context A compared to Context B for P24 (G, unpaired *t*-test: *t*_18_ = 2.46, *P* < 0.05) and P60 (H, unpaired *t*-test: *t*_18_ = 2.20, *P* < 0.05) mice. **(I to K)** Histograms depicting the distribution of angular positions of all time bins pooled across mice during the Context A and Context B tests for P20 (I), P24 (J), and P60 (K) mice. Number of time bins (Count) is shown along the radial axis, and the reward zone angular position (π/4) is highlighted in red. **(L to N)** During the test, the average angular position did not differ across Context A and Context B in P20 (L, unpaired *t*-test: *t*_18_ = 0.17, *P* = 0.83) or P24 (M, unpaired *t*-test: *t*_18_ = 0.18, *P* = 0.85) mice, but was more focused towards the previously rewarded position in Context A compared to Context B in P60 mice (N, unpaired *t*-test: *t*_18_ = 3.42, *P* < 0.01). **(O to Q)** Different groups of mice were trained in the spatial foraging task as before. Latencies to locate the reward zone decreased across trials (O) and did not differ across groups by the last training trial (P, ANOVA, no main effect of Age: *F*_2,27_ = 0.04, *P* = 0.95). When tested in Context A with the distal spatial cues removed, all groups performed at chance levels (Q, ANOVA, no main effect of Age: *F*_2,27_ = 0.07, *P* = 0.92). Data points are individual mice with mean ± s.e.m. * *P* < 0.05; ** *P* < 0.01; *** *P* < 0.001.

**Figure S3.**
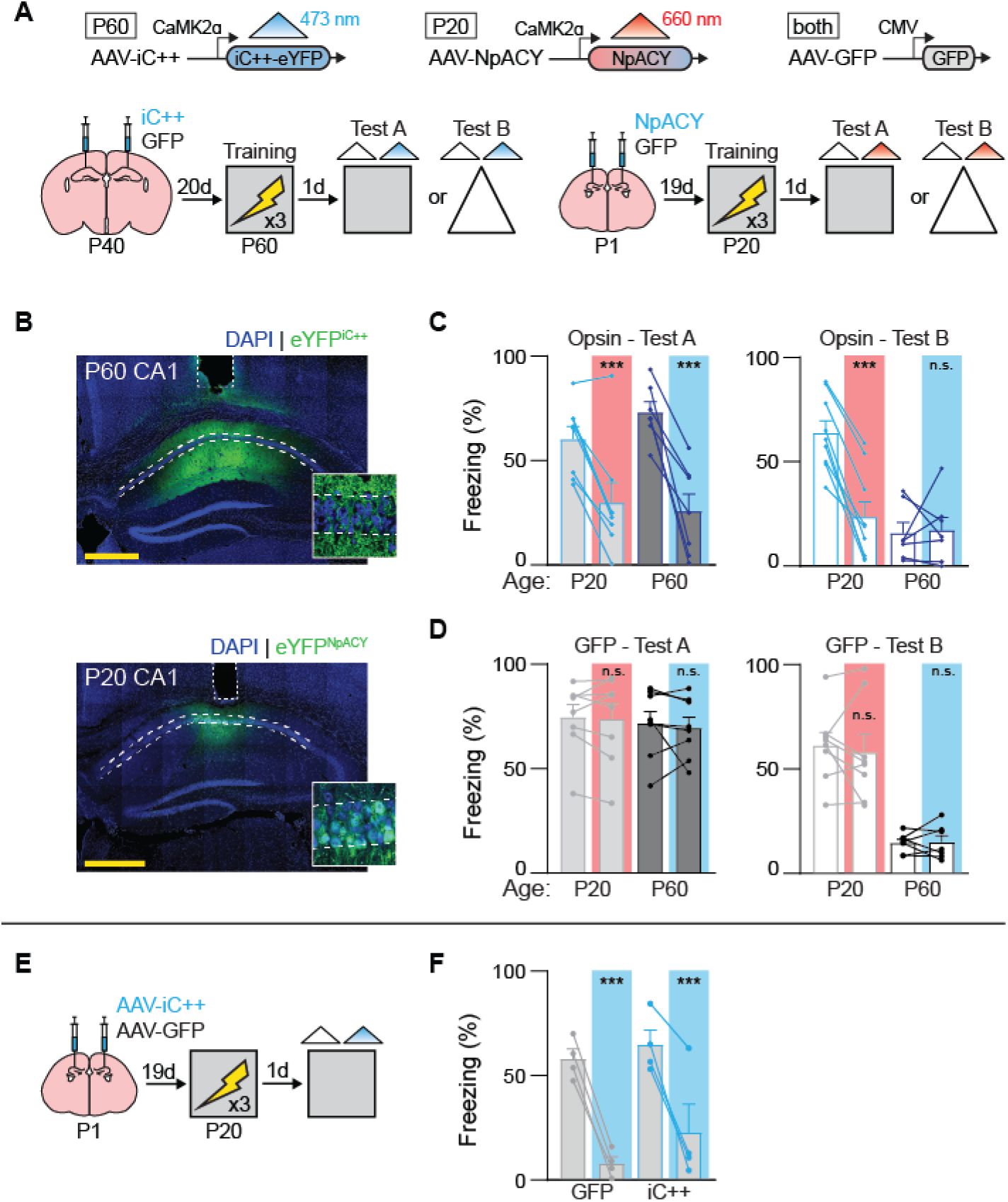
The CA1 is required for contextual fear memory recall in adult and juvenile mice. **(A)** Mice were microinjected with AAVs (P60: AAV-iC++ or AAV-GFP, P20: AAV-NpACY or AAV-GFP), trained in contextual fear conditioning, then tested the next day in Context A or B with no light followed by blue (iC++) or red (NpACY) light to silence CA1 pyramidal neurons. **(B)** Representative images showing iC++ and NpACY viral expression in CA1 of P60 and P20 mice. **(C)** Silencing CA1 pyramidal neurons during testing impaired memory retrieval for P20 and P60 mice in Context A (RM-ANOVA, no Age × Light interaction: *F*_1,13_ = 2.94, *P* = 0.10; no main effect of Age: *F*_1,13_ = 0.22, *P* = 0.64; main effect of Light: *F*_1,13_ = 62.74, *P* < 0.00001) and P20 mice in Context B (RM-ANOVA, Age × Light interaction: *F*_1,14_ = 28.80, *P* < 0.0001; main effect of Age: *F*_1,14_ = 11.43, *P* < 0.01; main effect of Light: *F*_1,14_ = 25.11, *P* < 0.001). **(D)** Red and blue light alone do not impair memory recall in P20 and P60 mice, respectively, in Context A (RM-ANOVA, no Age × Light interaction: *F*_1,14_ = 0.06, *P* = 0.80; no main effect of Age: *F*_1,14_ = 0.18, *P* = 0.67; no main effect of Light: *F*_1,14_ = 0.40, *P* = 0.53) or Context B (RM-ANOVA, no Age × Light interaction: *F*_1,13_ = 0.29, *P* = 0.59; main effect of Age: *F*_1,13_ = 34.14, *P* < 0.0001; no main effect of Light: *F*_1,13_ = 0.17, *P* = 0.68). **(E)** Mice were microinjected with AAV-iC++ or AAV-GFP, trained in contextual fear conditioning on P20, then tested the next day in Context A with no light followed by blue light to silence CA1 pyramidal neurons. **(F)** Blue light non-specifically reduced freezing behavior in P20 mice during the test (RM-ANOVA, no Virus × Light interaction: *F*_1,6_ = 0.92, *P* = 0.37; no main effect of Virus: *F*_1,6_ = 1.03, *P* = 0.34; main effect of Light: *F*_1,6_ = 118.38, *P* < 0.0001). Data points are individual mice with mean ± s.e.m. Scale bars: yellow = 500 *μ*m. *** *P* < 0.001.

**Figure S4.**
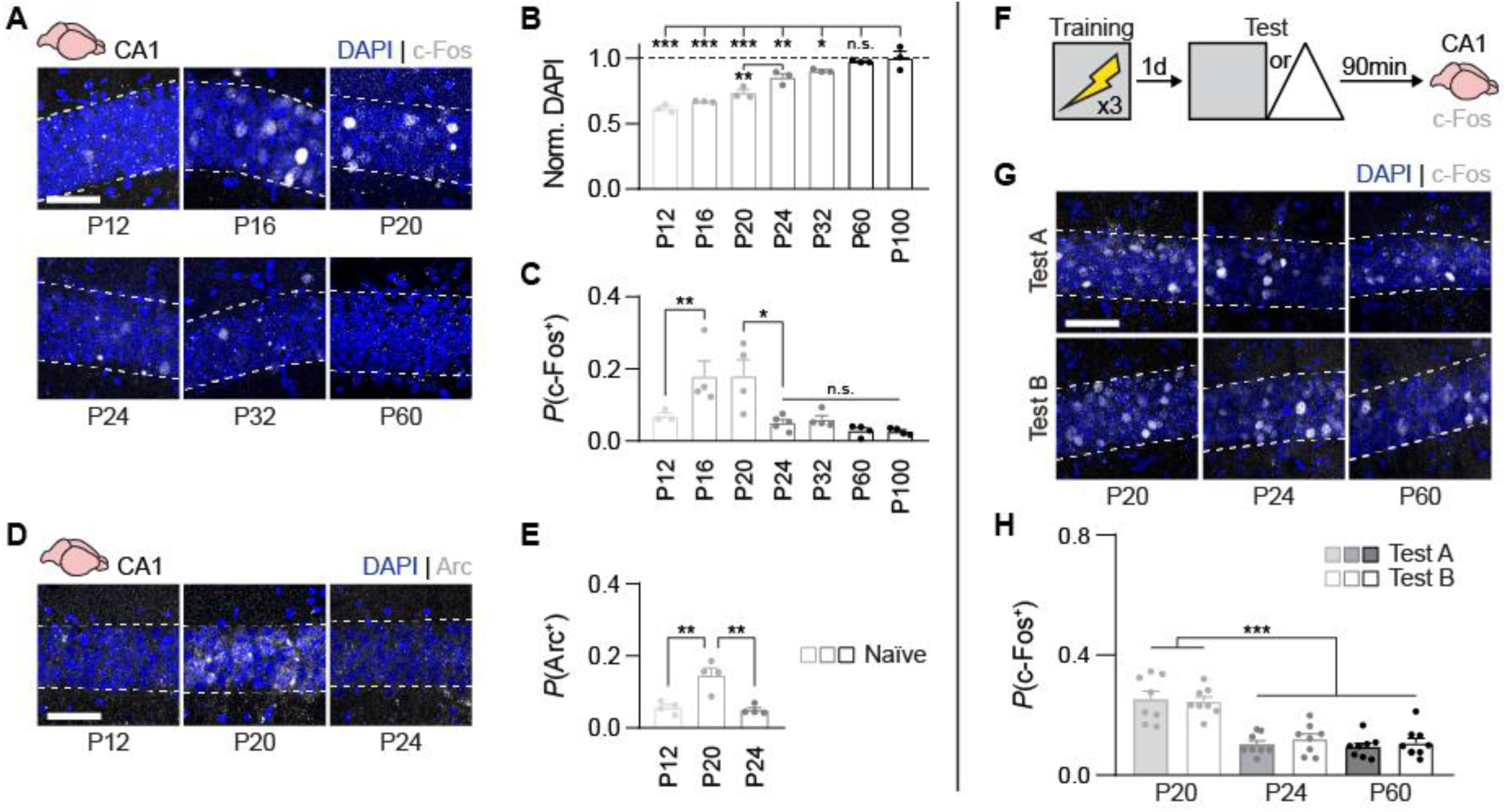
Immediate-early gene expression is transiently elevated in CA1 during the third postnatal week (Related to Figure 1). **(A)** Representative images showing c-Fos expression in CA1 pyramidal layer cells across development. **(B)** The density of DAPI^+^ cells in CA1 (normalized to the mean of mature adults, P100) increased into adulthood (ANOVA, main effect of Age: *F*_6,14_ = 34.42, *P* < 0.000001). **(C)** The proportion of pyramidal layer cells expressing c-Fos was elevated in naive ‘home cage’ mice during the third postnatal week (P16 to P20) (ANOVA, main effect of Age: *F*_6,21_ = 7.30, *P* < 0.001). **(D)** Representative images showing Arc expression in CA1 pyramidal layer cells across early development. **(E)** The proportion of pyramidal later cells expressing Arc was elevated in naive ‘home cage’ mice during the third postnatal week (P20) (ANOVA, main effect of Age: *F*_2,9_ = 15.75, *P* < 0.01). **(F)** Mice were trained in contextual fear conditioning, and c-Fos expression in dorsal CA1 was examined 90-min after the tests in Contexts A or B **(G)** Representative images showing c-Fos expression in CA1 pyramidal layer cells after contextual fear retrieval. **(H)** After testing, the proportion of pyramidal layer cells expressing c-Fos was elevated in P20 mice compared to P24 and P60 mice (ANOVA, no Age × Test Context interaction: *F*_2,43_ = 0.29, *P* = 0.74; main effect of Age: *F*_2,43_ = 43.30, *P* < 0.000001; no main effect of Test Context: *F*_1,43_ = 0.24, *P* = 0.62). Data points are individual mice with mean ± s.e.m. Scale bars: white = 50 *μ*m. * *P* < 0.05; ** *P* < 0.01; *** *P* < 0.001.

**Figure S5.**
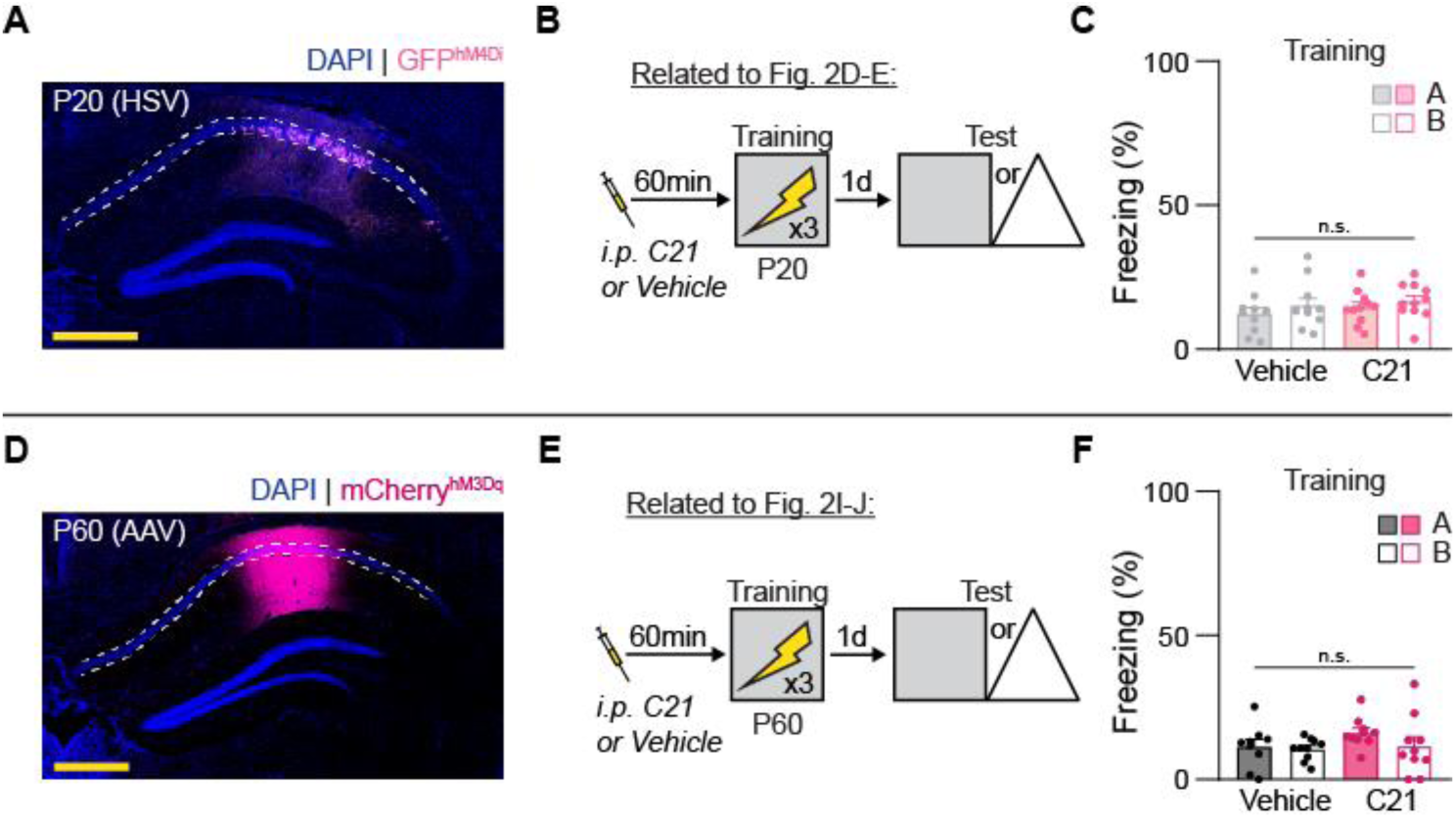
Chemogenetic manipulation of pyramidal neurons training data (Related to Figure 2). **(A)** Representative image showing hM4Di expression in dorsal CA1 of a P20 mouse. **(B to C)** Training data associated with Figure. 2D-E. Injecting P20 mice with C21 before training (B) did not alter freezing behavior during training (C, ANOVA, no Drug × Test Context interaction: *F*_1,40_ = 0.03, *P* = 0.85; no main effect of Drug: *F*_1,40_ = 0.72, *P* = 0.40; no main effect of Test Context: *F*_1,40_ = 1.46, *P* = 0.23). **(D)** Representative image showing hM3Dq expression in dorsal CA1 of a P60 mouse. **(E to F)** Training data associated with Figure 2I-J. Injecting P60 mice with C21 before training (E) did not alter freezing behavior during training (F, ANOVA, no Drug × Test Context interaction: *F*_1,35_ = 0.69, *P* = 0.40; no main effect of Drug: *F*_1,35_ = 1.66, *P* = 0.20; no main effect of Test Context: *F*_1,35_ = 1.48, *P* = 0.23). Data points are individual mice with mean ± s.e.m. Scale bars: yellow = 500 *μ*m.

**Figure S6.**
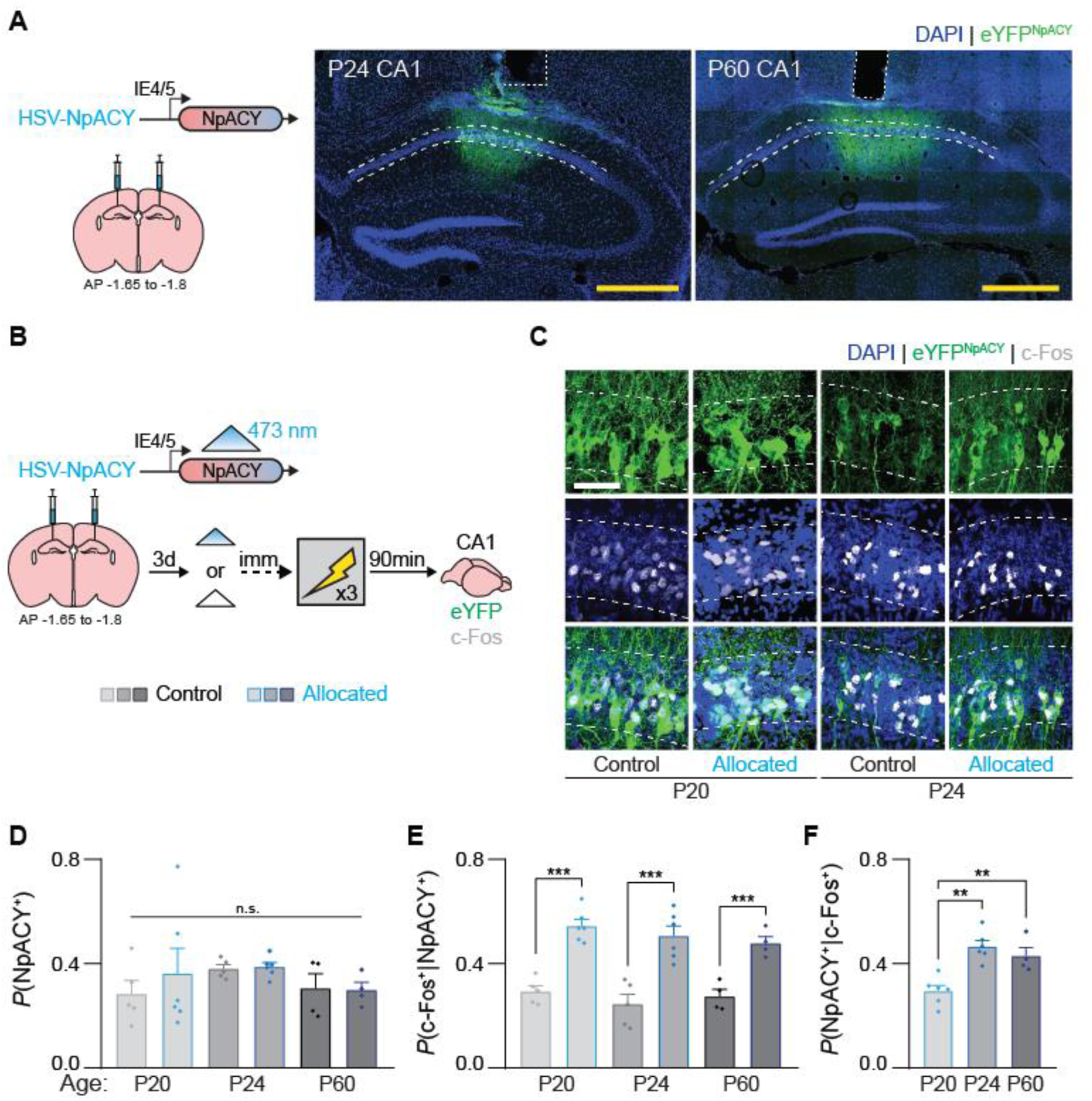
Optogenetics-mediated neuronal allocation is effective regardless of mouse age (Related to Figure 3). **(A)** Representative images showing HSV-NpACY expression in dorsal CA1 of P24 and P60 mice. **(B)** Mice were microinjected with HSV-NpACY in CA1 and three days later, blue light or no light was delivered for 30 s immediately before contextual fear conditioning training. c-Fos expression was examined 90-min after training. **(C)** Representative images showing NpACY and c-Fos expression in dorsal CA1. **(D to F)** The proportion of neurons expressing HSV-NpACY did not differ across groups (D, ANOVA, no Age × Light interaction: *F*_2,24_ = 0.72, *P* = 0.72; no main effect of Age: *F*_2,24_ = 1.09, *P* = 0.35; no main effect of Light: *F*_1,24_ = 0.31, *P* = 0.57). Blue-light stimulation immediately before training resulted in greater c-Fos expression in NpACY^+^ neurons in all age groups (E, ANOVA, no Age × Light interaction: *F*_2,24_ = 0.45, *P* = 0.63; no main effect of Age: *F*_2,24_ = 1.35, *P* = 0.27; main effect of Light: *F*_1,24_ = 86.88, *P* < 0.000001), but c-Fos was expressed in more NpACY^-^ neurons in P20 mice compared with P24 and P60 mice (F, ANOVA, main effect of Age: *F*_2,13_ = 14.17, *P* < 0.001). Data points are individual mice with mean ± s.e.m. Scale bars: white = 50 *μ*m, yellow = 500 *μ*m. * *P* < 0.05; ** *P* < 0.01; *** *P* < 0.001.

**Figure S7.**
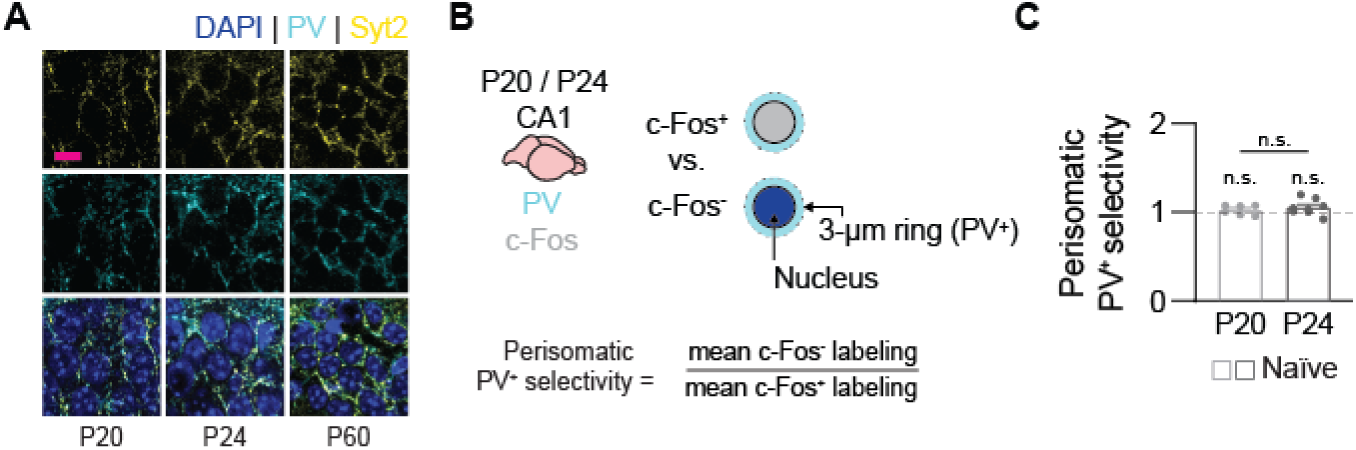
Perisomatic PV^+^ selectivity in experimentally naive mice (Related to Figure 4). **(A)** Representative images showing co-localization of Syt2^+^ synaptic puncta among PV^+^ neurites across development. **(B)** Perisomatic PV^+^ selectivity analysis was performed on CA1 pyramidal layer cells from naive P20 and P24 mice as a negative control. **(C)** PV^+^ neurites were not selectively localized around c-Fos^-^ compared to c-Fos^+^ cells in naive P20 and P24 mice (one-sample *t*-tests [with Bonferroni correction, ɑ = 0.025], P20: *t*_5_ = 1.25, *P* = 0.26; P24: *t*_7_ = 1.84, *P* = 0.10; P60: *t*_5_ = 5.85, *P* < 0.01; unpaired *t*-test P20 vs. P24: *t*_12_ = 0.72, *P* = 0.48). Data points are individual mice with mean ± s.e.m. Scale bars: magenta = 10 *μ*m.

**Figure S8.**
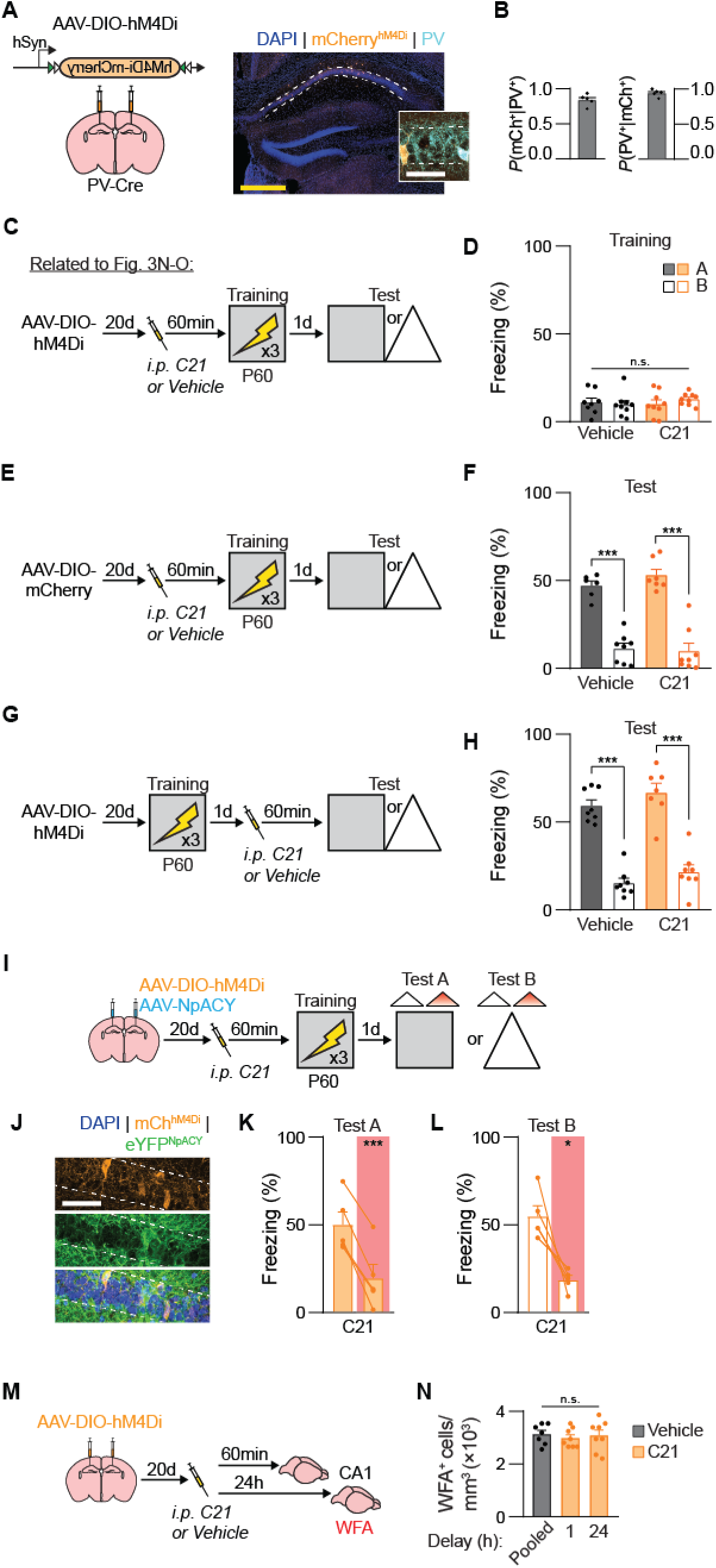
Control experiments for using chemogenetics to inhibit CA1 PV^+^ interneurons (Related to Figure 4). **(A to B)** AAV-DIO-hM4Di was expressed in dorsal CA1 (A) resulting in high penetrance (left) and specificity (right) of hM4Di in PV^+^ cells (B). **(C to D)** Training data associated with Figure 4N-O. Injecting P60 PV::hM4Di mice with C21 before training (C) did not alter freezing behavior during training (D, ANOVA, no Drug × Test Context interaction: *F*_1,31_ = 1.04, *P* = 0.31; no main effect of Drug: *F*_1,31_ = 0.26, *P* = 0.60; no main effect of Test Context: *F*_1,31_ = 0.11, *P* = 0.73). **(E to F)** Injecting P60 PV::mCherry mice with C21 before training (E) did not alter memory precision (F, ANOVA, no Drug × Test Context interaction: *F*_1,25_ = 1.07, *P* = 0.30; no main effect of Drug: *F*_1,25_ = 0.42, *P* = 0.51; main effect of Test Context: *F*_1,25_ = 123.31, *P* < 0.000001). **(G to H)** Injecting PV::hM4Di mice with C21 before testing (G) did not alter memory precision (H, ANOVA, no Drug × Test Context interaction: *F*_1,27_ = 0.019, *P* = 0.88; no main effect of Drug: *F*_1,27_ = 3.22, *P* = 0.08; main effect of Test Context: *F*_1,27_ = 134.56, *P* < 0.000001). **(I to L)** Mice were microinjected with AAV-DIO-hM4Di and AAV-NpACY in dorsal CA1, injected with C21 1-h before training, then tested the next day in Context A or Context B without and with red light to silence CA1 pyramidal neurons (I). Representative images of CA1 showing hM4Di expression in PV^+^ interneurons and NpACY in excitatory neurons (J). Inhibiting CA1 PV^+^ interneurons before training in P60 mice resulted in imprecise memories that were still dependent on the hippocampus, as silencing CA1 neurons with red light reduced freezing in Context A (K, paired *t*-test: *t*_4_ = 9.77, *P* < 0.001) and in Context B (L, paired *t*-test: *t*_4_ = 4.30, *P* < 0.05). **(M to N)** Adult PV-Cre mice were microinjected with AAV-DIO-hM4Di in dorsal CA1, injected with C21, and WFA expression in dorsal CA1 was examined 1 or 24 h later(M). Acute PV^+^ interneuron inhibition did not alter WFA^+^ PNN density at timepoints relevant to contextual fear conditioning training or testing (N, ANOVA, no main effect of Drug/Delay: *F*_2,20_ = 0.20, *P* = 0.81). Data points are individual mice with mean ± s.e.m. Scale bars: white = 50 *μ*m, yellow = 500 *μ*m. * *P* < 0.05; ** *P* < 0.01; *** *P* < 0.001.

**Figure S9.**
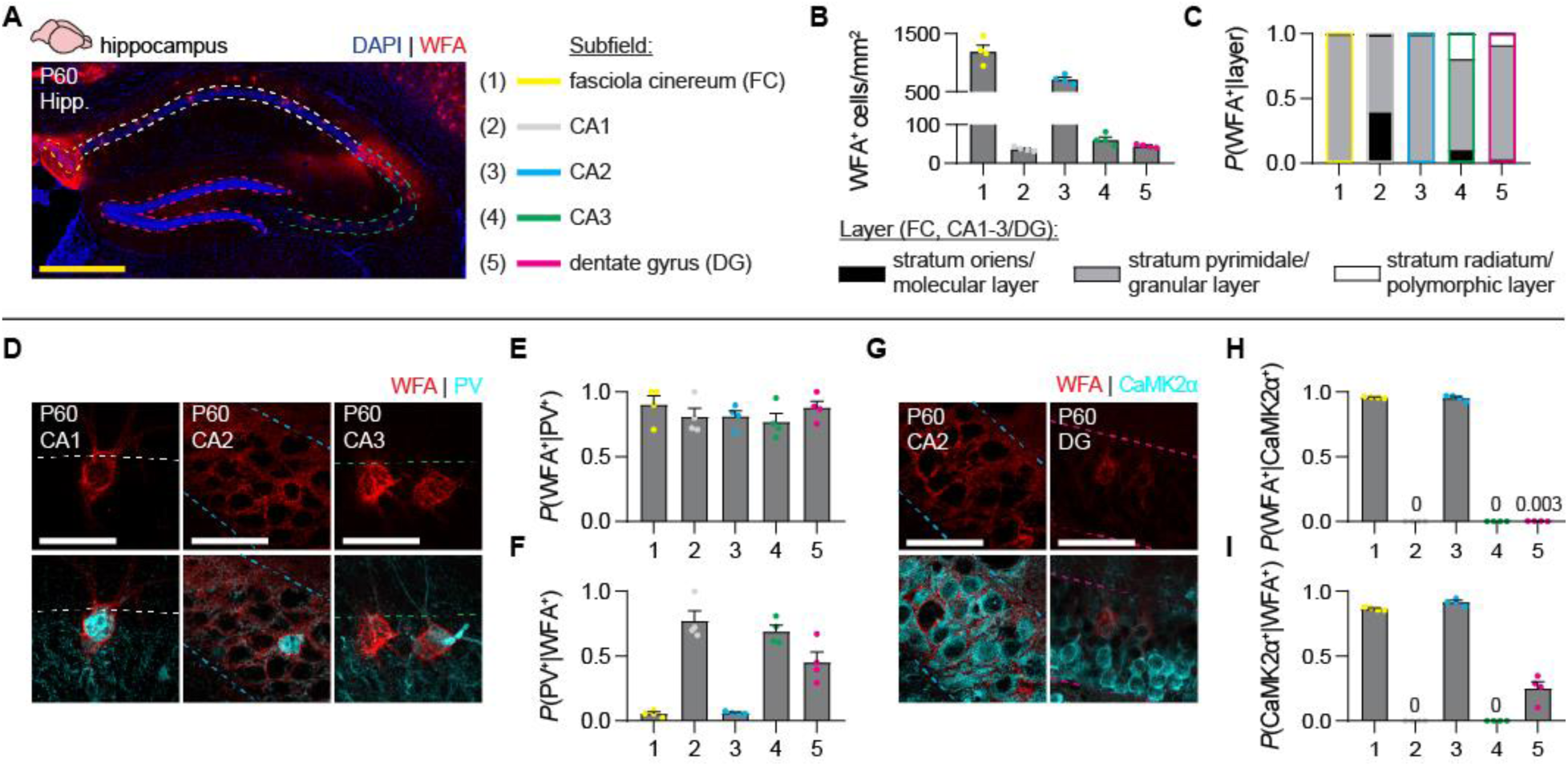
Characterization of PNNs in the adult mouse dorsal hippocampus. **(A)** PNNs were identified in the P60 dorsal hippocampus using WFA staining. **(B)** WFA^+^ PNNs were sparsely distributed in subfields CA1, CA3, and dentate gyrus (DG), and densely packed in fasciola cinereum (FC) and CA2. **(C)** Across all subfields, most WFA^+^ PNNs were located in the pyramidal/granule cell layer. **(D)** Example images showing colocalization of WFA^+^ PNNs with PV^+^ interneurons in different subfields. **(E)** The majority PV^+^ interneurons in the dorsal hippocampus were surrounded by WFA^+^ PNNs. **(F)** A minority WFA^+^ PNNs surrounded PV^-^ cells in CA1 and CA3, and to a lesser extent in DG. **(G)** Example images showing colocalization of WFA^+^ PNNs with CaMK2ɑ^+^ excitatory neurons in different subfields. **(H)** The majority CaMK2ɑ^+^ neurons in FC and CA2 were surrounded by WFA^+^ PNNs, and no CaMK2ɑ^+^ neurons in CA1 and CA3 were surrounded by WFA^+^ PNNs. A small proportion of CaMK2ɑ^+^ neurons (likely, mature granule cells based on proximity to the hilus) were surrounded by WFA^+^ PNNs. **(I)** A minority WFA^+^ PNNs surrounded CaMK2ɑ^-^ cells in FC and CA2. Data points are individual mice with mean ± s.e.m. Scale bars: white = 50 *μ*m, yellow = 500 *μ*m.

**Figure S10.**
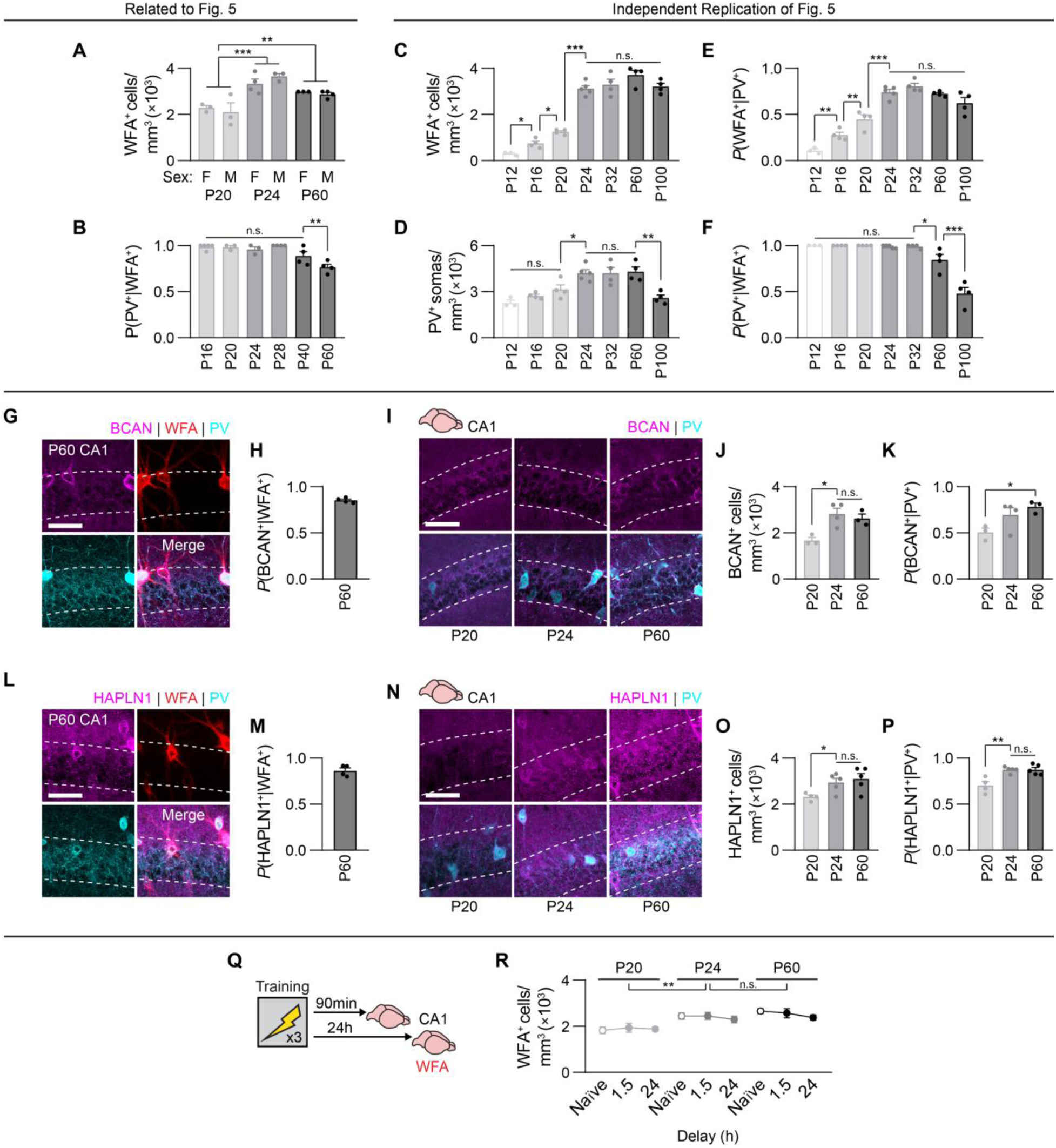
Maturation of CA1 PNNs (Related to Figure 5). **(A)** The maturation of CA1 PNNs by P24 was not sex-dependent (ANOVA, no Age × Sex interaction: *F*_2,14_ = 1.03, *P* = 0.38; main effect of Age: *F*_2,14_ = 22.13, *P* < 0.0001; no main effect of Sex: *F*_1,14_ = 0.004, *P* = 0.95). **(B)** A small proportion of WFA^+^ PNNs surrounding PV^-^ cells form in CA1 during young adulthood (ANOVA, main effect of Age: *F*_5,17_ = 10.94, *P* < 0.0001). **(C to F)** Independent replication of Figure 5C-D with expanded Age range. The density of WFA^+^ PNNs (C, ANOVA, main effect of Age: *F*_6,21_ = 82.50, *P* < 0.000001), PV^+^ interneurons (D, ANOVA, main effect of Age: *F*_6,21_ = 10.70, *P* < 0.0001), and proportion of PV^+^ interneurons surrounded by WFA^+^ PNNs (E, ANOVA, main effect of Age: *F*_6,21_ = 46.68, *P* < 0.000001) reach adult-like levels by P24, with WFA^+^PV^-^ PNNs developing during adulthood (F, ANOVA, main effect of Age: *F*_6,21_ = 32.30, *P* < 0.000001). **(G to H)** Representative images (G) showing high colocalization of BCAN with WFA^+^ PNNs in P60 CA1 (H). **(I to K)** Representative images (I) showing increased density of BCAN^+^ cells (J, ANOVA, main effect of Age: *F*_2,7_ = 8.40, *P* < 0.05) and colocalization of BCAN and PV^+^ interneurons (K, ANOVA, main effect of Age: *F*_2,7_ = 4.70, *P* = 0.05) across development in dorsal CA1. **(L to M)** Representative images (M) showing high colocalization of HAPLN1 with WFA^+^ PNNs in P60 CA1 (M). **(N to P)** Representative images (N) showing increased density of HAPLN1^+^ cells (O, ANOVA, main effect of Age: *F*_2,11_ = 4.55, *P* < 0.05) and colocalization of BCAN and PV^+^ interneurons (P, ANOVA, main effect of Age: *F*_2,11_ = 12.05, *P* < 0.01) across development in dorsal CA1. **(Q to R)** WFA expression in dorsal CA1 was examined 90-min or 24-h after contextual fear conditioning (Q). Training did not alter the trajectory of WFA^+^ PNN development in CA1 (R, ANOVA, no Age × Delay interaction: *F*_4,52_ = 1.92, *P* = 0.12; main effect of Age: *F*_2,52_ = 5.88, *P* < 0.01; no main effect of Delay: *F*_2,52_ = 0.79, *P* = 0.45). Data points are individual mice with mean ± s.e.m. Scale bars: white = 50 *μ*m. * *P* < 0.05; ** *P* < 0.01; *** *P* < 0.001

**Figure S11.**
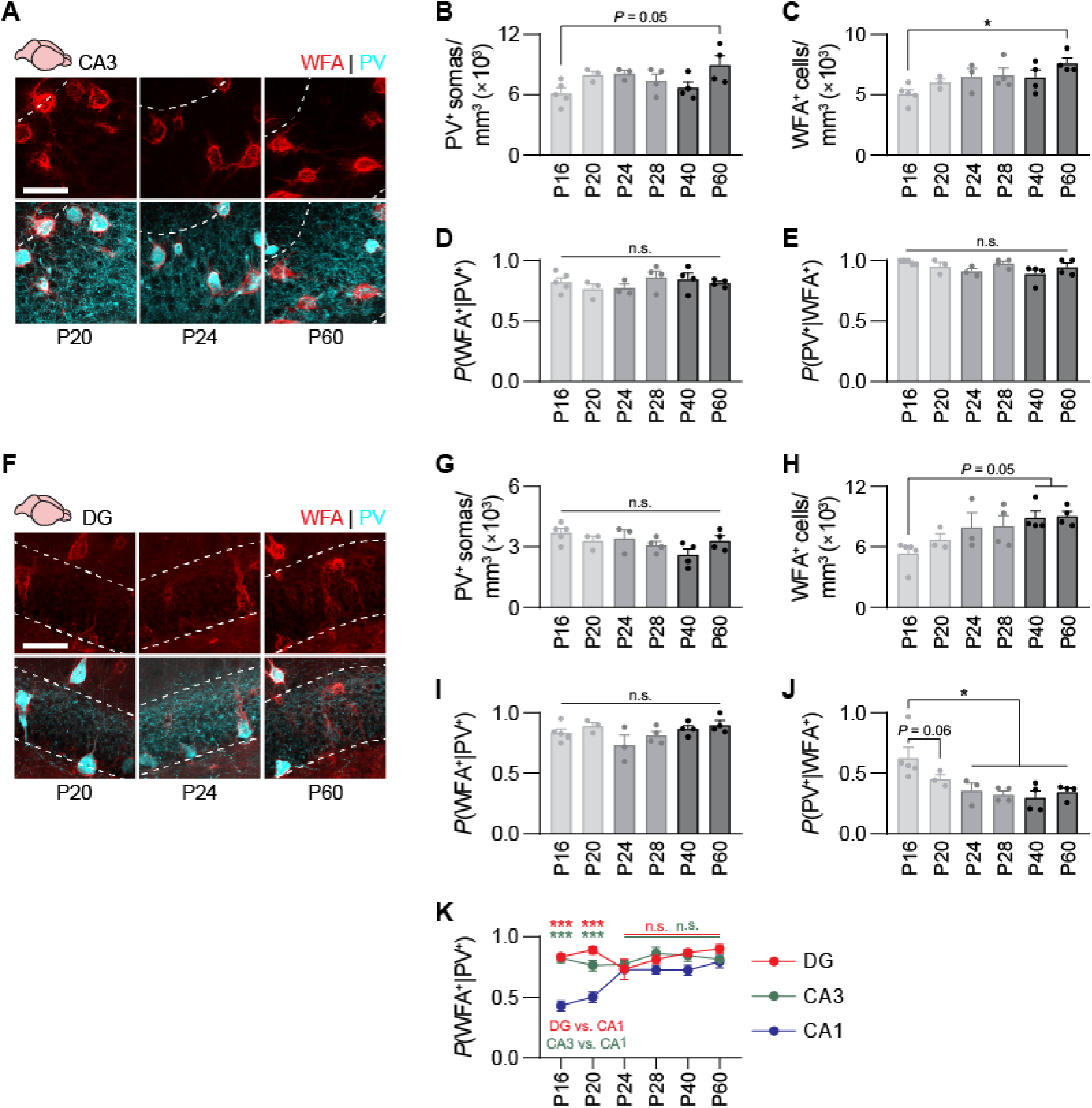
Maturation of CA3 and DG PNNs (Related to Figure 5). **(A)** Representative images showing WFA^+^ PNNs surrounding PV^+^ interneurons in CA3 across development. **(B to E)** In CA3, he density of PV^+^ interneurons (B, ANOVA, main effect of Age: *F*_5,17_ = 2.87, *P* < 0.05) and WFA^+^ PNNs (C, ANOVA, main effect of Age: *F*_5,17_ = 3.17, *P* < 0.05) increased slightly between P16 and P60. The proportion of PV^+^ interneurons surrounded by WFA^+^ PNNs reached adult-like levels before P16 (D, ANOVA, main effect of Age: *F*_5,17_ = 0.87, *P* = 0.51), and no further development of WFA^+^ PNNs around PV^-^ cells from P16 to P60 (E, ANOVA, main effect of Age: *F*_5,17_ = 2.18, *P* = 0.10). **(F)** Representative images showing WFA^+^ PNNs surrounding PV^+^ interneurons in DG across development. **(G to J)** In DG, the density of PV^+^ interneurons did not change (G, ANOVA, main effect of Age: *F*_5,17_ = 2.34, *P* = 0.08) and WFA^+^ PNNs increased (H, ANOVA, main effect of Age: *F*_5,17_ = 3.46, *P* < 0.05) between P16 and P60. The proportion of PV^+^ interneurons surrounded by WFA^+^ PNNs reached adult-like levels before P16 (I, ANOVA, main effect of Age: *F*_5,17_ = 2.02, *P* = 0.12), and WFA^+^ PNNs around PV^-^ cells increased from P16 to P60 (E, ANOVA, main effect of Age: *F*_5,17_ = 4.85, *P* < 0.01). **(K)** Development of WFA^+^ PNNs around PV^+^ interneurons was delayed in CA1 compared to CA3 and DG (RM-ANOVA, Age × Region interaction: *F*_10,34_ = 7.42, *P* < 0.00001; main effect of Age: *F*_5,17_ = 4.05, *P* < 0.05; main effect of Region: *F*_2,34_ = 48.82, *P* < 0.000001). Data shown in Panel S11K are re-plotted from Fig. 4D, S11D, and S11I, and re-analyzed. Data points are individual mice with mean ± s.e.m. Scale bars: white = 50 *μ*m. * *P* < 0.05; ** *P* < 0.01; *** *P* < 0.001.

**Figure S12.**
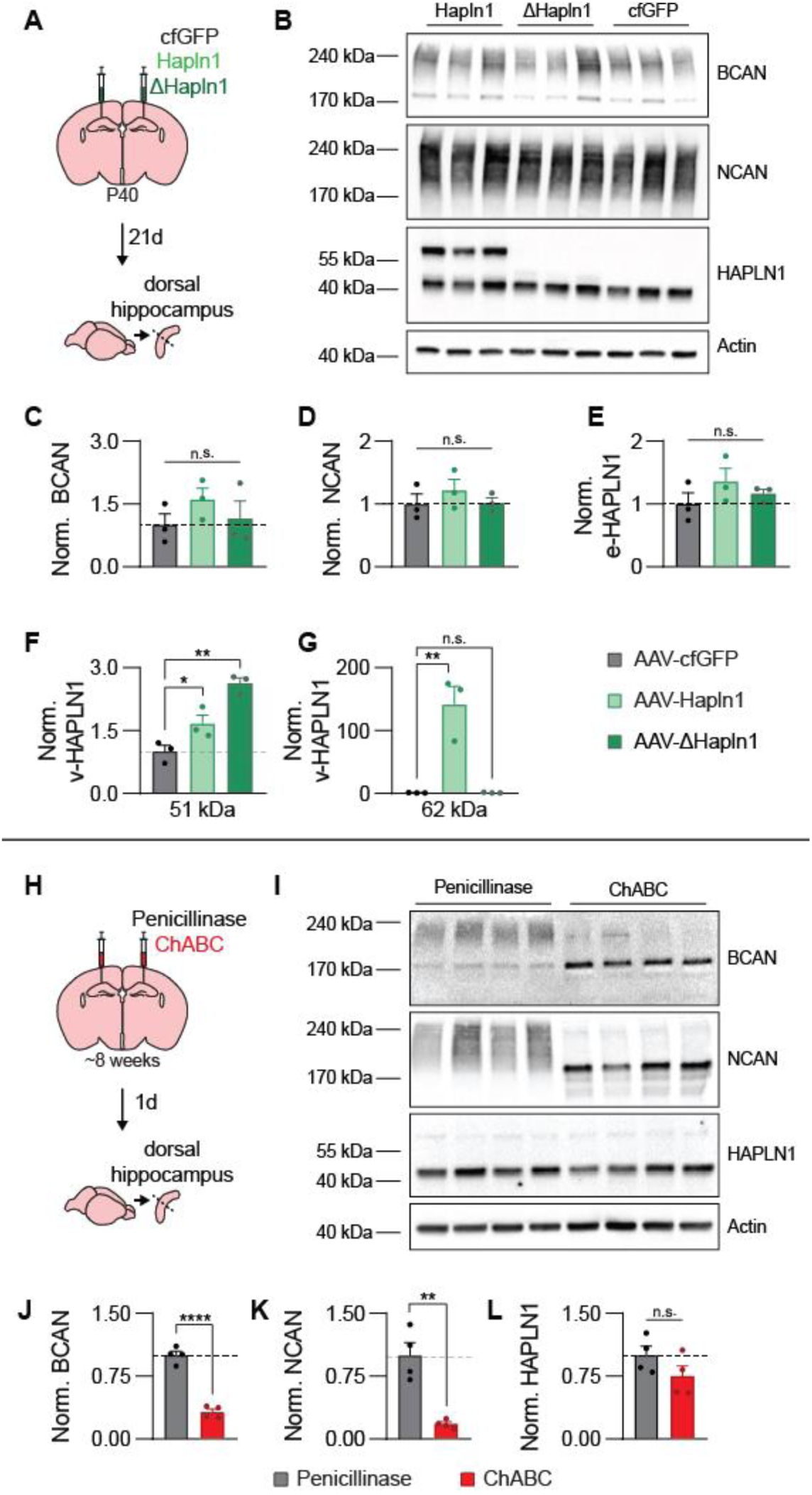
Viral constructs targeting HAPLN1 do not alter endogenous CSPG or HAPLN1 expression (Related to Figure 5). **(A)** Mice were microinjected with AAV-cfGFP, AAV-Hapln1, or AAV-ΔHapln1 in the dorsal hippocampus. Three weeks later, hippocampi of adult mice were dissected and used for western blot analyses. **(B)** Western blot illustrating changes in endogenous brevican (BCAN, 145 kDa), neurocan (NCAN, 130 kDa), HAPLN1 (e-HAPLN1, 41 kDa), and virally expressed HAPLN1 (v-HAPLN1, 51 and 62 kDa) in dorsal hippocampus extracts following AAV expression. Actin (42 kDa) was used as a loading control. Individual lanes are different mice. **(C to G)** AAV-Hapln1 and AAV-ΔHapln1 did not alter BCAN (C, ANOVA, no main effect of Virus: *F*_2,6_ = 0.69, *P* = 0.53), NCAN (D, ANOVA, no main effect of Virus: *F*_2,6_ = 0.92, *P* = 0.44), or e-HAPLN1 (E, ANOVA, no main effect of Virus: *F*_2,6_ = 1.18, *P* = 0.36) expression, but increased v-HAPLN1 expression (F, 51 kDa: ANOVA, main effect of Virus: *F*_2,6_ = 24.71, *P* < 0.01; G, 62 kDa: ANOVA, main effect of Virus: *F*_2,6_ = 23.40, *P* < 0.01). **(H)** Mice were microinjected with Penicillinase or ChABC in the dorsal hippocampus. One day later, hippocampi of adult mice were dissected and used for western blot analyses. **(I)** Western blot illustrating changes in endogenous BCAN, NCAN, HAPLN1 in dorsal hippocampus extracts following enzyme injection. Actin was used as a loading control. Individual lanes are different mice. **(J to L)** ChABC treatment reduced BCAN (J, unpaired *t*-test: *t*_6_ = 11.23, *P* < 0.0001), NCAN (K, unpaired *t*-test: *t*_6_ = 5.41, *P* < 0.01), but not HAPLN1 (L, unpaired *t*-test: *t*_6_ = 1.53, *P* = 0.17) in dorsal hippocampus. Data points are individual mice with mean ± s.e.m. Data are normalized to the mean of the control group (AAV-cfGFP or Penicillinase). * *P* < 0.05; ** *P* < 0.01; *** *P* < 0.001; **** *P* < 0.0001.

**Figure S13.**
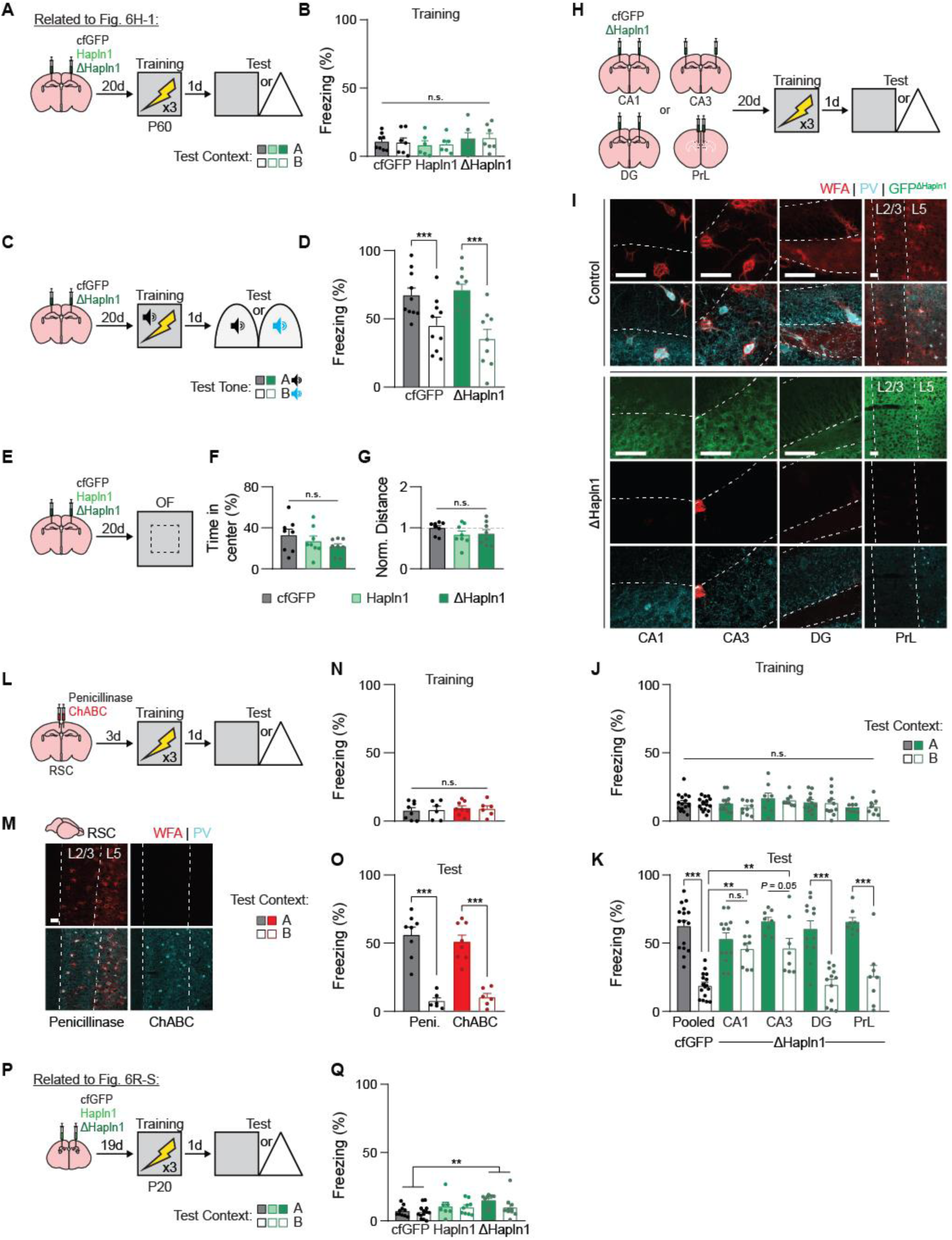
Hippocampal (and not cortical) PNNs are necessary for contextual memory precision in adult mice (Related to Figure 6). **(A to B)** Training data associated with Figure 6H-I. Expression of AAV-Hapln1 or AAV-ΔHapln1 in the dorsal CA1 (A) did not alter P60 mouse freezing behavior during training (B, ANOVA, no Virus × Test Context interaction: *F*_2,34_ = 0.022, *P* = 0.97; no main effect of Virus: *F*_2.34_ = 1.00, *P* = 0.37; no main effect of Test Context: *F*_1,34_ = 0.0034, *P* = 0.96). **(C to D)** Expression of AAV-ΔHapln1 in the dorsal CA1 before tone fear conditioning (C) did not alter memory precision for similar auditory cues (D, ANOVA, no Virus × Test Tone interaction: *F*_1,35_ = 1.28, *P* = 0.26; no main effect of Virus: *F*_1,35_ = 0.28, *P* = 0.59; main effect of Test Tone: *F*_1,35_ = 24.68, *P* < 0.0001). **(E to G)** Expression of AAV-Hapln1 or AAV-ΔHapln1 in the dorsal CA1 before an open field test (E) did not alter anxiety-like behavior (F, ANOVA, no main effect of Virus: *F*_2,21_ = 1.43, *P* = 0.26) or locomotion (G, ANOVA, no main effect of Virus: *F*_2,21_ = 1.14, *P* = 0.33) in adult mice. **(H to K)** Expression of AAV-ΔHapln1 in CA1, CA3, or DG of the hippocampus, or the prelimbic region (PrL) of the medial prefrontal cortex before contextual fear conditioning (H) destabilized PNNs in the corresponding brain region (I), but did not alter freezing behavior during training (J, ANOVA, no Virus/Region × Test Context interaction: *F*_4,100_ = 0.20, *P* = 0.93; no main effect of Virus/Region: *F*_4,100_ = 1.86, *P* = 0.12; no main effect of Test Context: *F*_1,100_ = 0.86, *P* = 0.35). Destabilizing PNNs in CA1 or CA3 (but not DG or PrL) with AAV-ΔHapln1 reduced memory precision in adult mice during the memory test (K, ANOVA, Virus/Region × Test Context interaction: *F*_4,100_ = 5.84, *P* < 0.001; main effect of Virus/Region: *F*_4,100_ = 3.67, *P* < 0.01; main effect of Test Context: *F*_1,100_ = 95.72, *P* < 0.000001). **(L to O)** Microinjection of ChABC into the retrosplenial cortex (RSC) before contextual fear conditioning (L) digested PNNs (M), and did not alter freezing behavior during training (N, ANOVA, no Microinjection × Test Context interaction: *F*_1,24_ = 0.05, *P* = 0.82; no main effect of Microinjection: *F*_1,24_ = 0.33, *P* = 0.56; no main effect of Test Context: *F*_1,24_ = 0.0015, *P* = 0.96) or during the memory test (O, ANOVA, no Microinjection × Test Context interaction: *F*_1,24_ = 0.75, *P* = 0.39; no main effect of Microinjection: *F*_1,24_ = 0.06, *P* = 0.79; main effect of Test Context: *F*_1,24_ = 92.64, *P* < 0.000001). **(P to Q)** Training data associated with Figure 6R-S. Expression of AAV-Hapln1 in the dorsal CA1 (P) did not alter P20 mouse freezing behavior during training (Q, ANOVA, no Virus × Test Context interaction: *F*_2,53_ = 0.98, *P* = 0.37; main effect of Virus: *F*_2,53_ = 5.17, *P* < 0.01; no main effect of Test Context: *F*_1,53_ = 1.96, *P* = 0.16). Data points are individual mice with mean ± s.e.m. Scale bars: white = 50 *μ*m. * *P* < 0.05; ** *P* < 0.01; *** *P* < 0.001.

**Figure S14.**
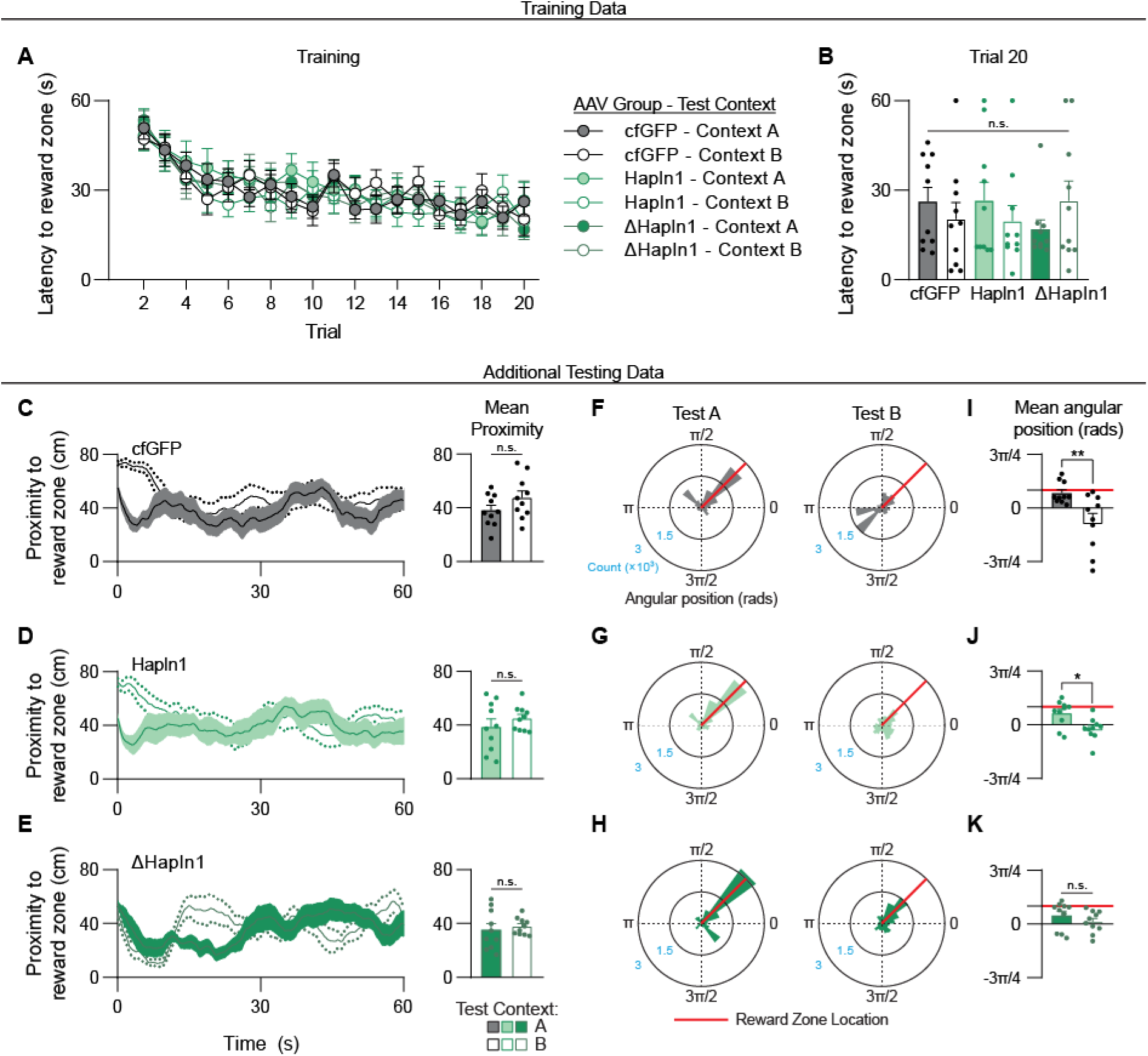
Spatial foraging task training data and additional test performance measures in adult mice with viral PNN manipulations (Related to Figure 6). **(A to B)** Latencies to locate the reward zone decreased across training trials in all groups (A) and did not differ across groups by the last training trial (B, ANOVA, no Virus × Test Context interaction: *F*_2,54_ = 1.45, *P* = 0.24; no main effect of Virus: *F*_2,54_ = 0.05, *P* = 0.94; no main effect of Test Context: *F*_1,54_ = 0.05, *P* = 0.81). **(C to E)** During the test, the average proximity to the previously rewarded zone in Context A or the equivalent area in Context B did not differ in AAV-cfGFP (C, unpaired *t*-test: *t*_18_ = 1.48, *P* = 0.15), AAV-Hapln1 (D, unpaired *t*-test: *t*_18_ = 0.95, *P* = 0.35), or AAV-ΔHapln1 (E, unpaired *t*-test: *t*_18_ = 0.48, *P* = 0.63) mice. **(F to H)** Histograms depicting the distribution of angular positions of all time bins pooled across mice during the Context A and Context B tests for AAV-cfGFP (F), AAV-Hapln1 (G), and AAV-ΔHapln1 (H) mice. Number of time bins (Count) is shown along the radial axis, and the reward zone angular position (π/4) is highlighted in red. **(I to K)** During the test, the average angular position of mice was more focused towards the previously rewarded position in Context A compared to Context B in AAV-cfGFP (I, unpaired *t*-test: *t*_18_ = 3.01, *P* < 0.01) and AAV-Hapln1 (J, unpaired *t*-test: *t*_18_ = 2.87, *P* < 0.05) groups, but not the AAV-ΔHapln1 (K, unpaired *t*-test: *t*_18_ = 1.08, *P* = 0.29) group. Data points are individual mice with mean ± s.e.m. * *P* < 0.05; ** *P* < 0.01.

**Figure S15.**
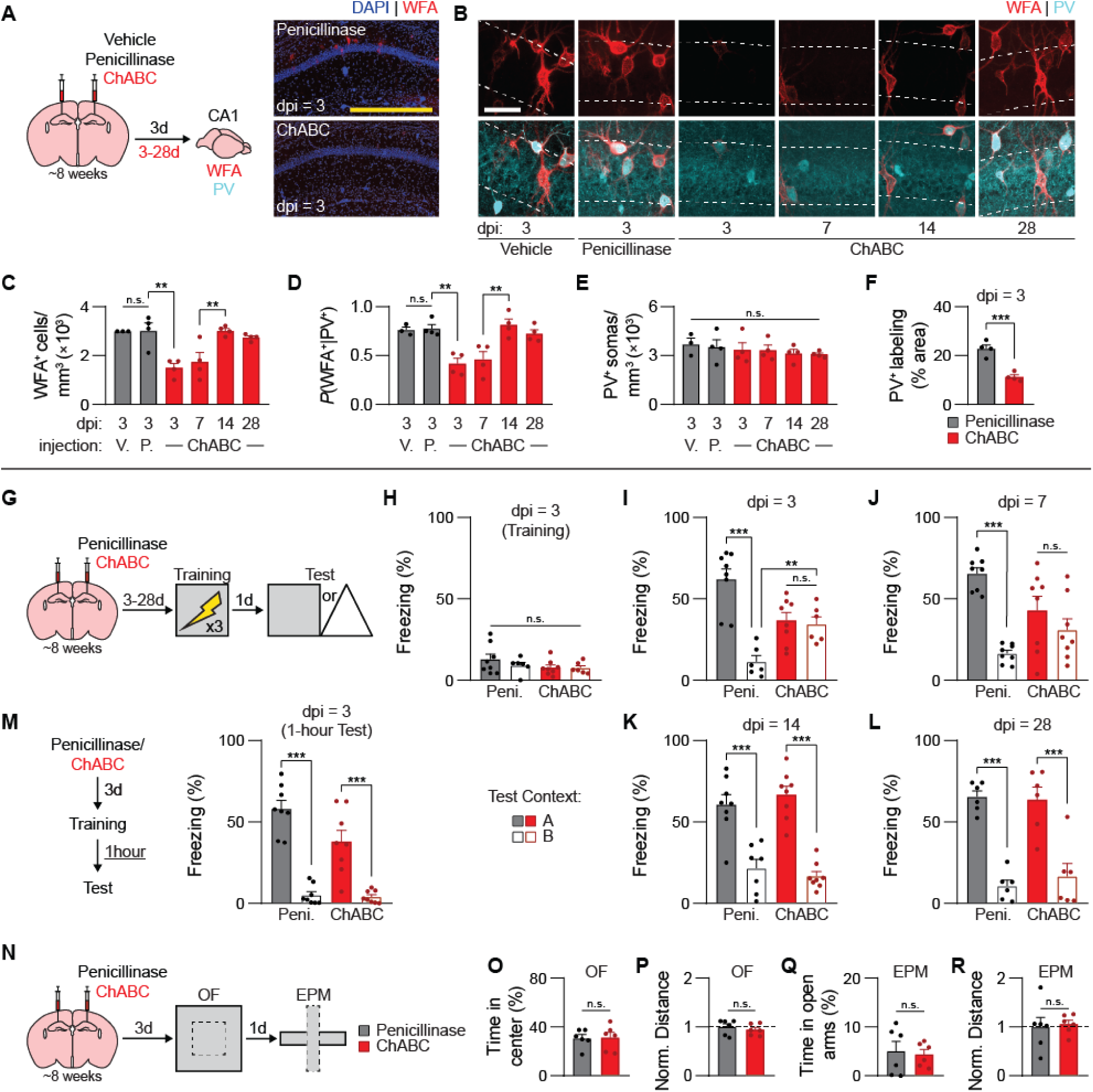
Enzymatic digestion of CA1 PNNs with ChABC reinstates juvenile-like memory imprecision. **(A)** Chondroitinase ABC (ChABC) was microinjected into the dorsal CA1 of adult mice to rapidly digest PNNs, and mice were perfused 3-28 d post-injection (dpi). **(B)** Representative images showing WFA^+^ PNNs and PV^+^ interneurons in dorsal CA1 at different dpi. **(C to F)** ChABC treatment transiently reduced the density of WFA^+^ PNNs (C, ANOVA, main effect of Injection/Delay: *F*_5,17_ = 9.14, *P* < 0.001) surrounding PV^+^ interneurons (D, ANOVA, main effect of Injection/Delay: *F*_5,17_ = 9.58, *P* < 0.001) at 3-7dpi, with eventual regeneration of PNNs occurring by 14-28 dpi. ChABC treatment did not alter PV^+^ interneuron density (E, ANOVA, main effect of Injection/Delay: *F*_5,17_ = 0.41, *P* = 0.83), but loss of PNNs led to a retraction of PV^+^ neurites in the pyramidal cell layer (F, unpaired *t*-test: *t*_6_ = 6.37, *P* < 0.001). **(G to L)** Adult mice were microinjected with Penicillinase or ChABC at different time points before contextual fear conditioning (G). ChABC treatment did not alter freezing behavior during training (H, ANOVA, no Injection × Test Context interaction: *F*_1,24_ = 0.55, *P* = 0.46; no main effect of Injection: *F*_1,24_ = 1.86, *P* = 0.18; no main effect of Test Context: *F*_1,24_ = 0.69, *P* = 0.41) but reduced memory precision at and 3- (I, ANOVA, Injection × Test Context interaction: *F*_1,24_ = 20.40, *P* < 0.001; no main effect of Injection: *F*_1,24_ = 0.04, *P* = 0.82; main effect of Test Context: *F*_1,24_ = 24.41, *P* < 0.0001) and 7-dpi (J, ANOVA, Injection × Test Context interaction: *F*_1,28_ = 9.64, *P* < 0.01; no main effect of Injection: *F*_1,28_ = 0.46, *P* = 0.49; main effect of Test Context: *F*_1,28_ = 26.27, *P* < 0.0001) when PNNs levels remained low. Adult-like memory precision was restored once PNN levels recovered at 14- (K, ANOVA, no Injection × Test Context interaction: *F*_1,27_ = 1.13, *P* = 0.29; no main effect of Injection: *F*_1,27_ = 0.019, *P* = 0.89; main effect of Test Context: *F*_1,27_ = 74.82, *P* < 0.000001) and 28-dpi (L, ANOVA, no Injection × Test Context interaction: *F*_1,20_ = 0.37, *P* = 0.54; no main effect of Injection: *F*_1,20_ = 0.10, *P* = 0.74; main effect of Test Context: *F*_1,20_ = 68.26, *P* < 0.000001). **(M)** PNN digestion with ChABC did not reduce precision for short-term (1 h) memories (ANOVA, Injection × Test Context interaction: *F*_1,28_ = 4.41, *P* < 0.05; main effect of Injection: *F*_1,28_ = 5.46, *P* < 0.05; main effect of Test Context: *F*_1,28_ = 90.74, *P* < 0.000001). **(N to R)** Microinjection of ChABC into dorsal CA1 of adult mice before an open field and elevated-plus maze exploration (N), did not alter anxiety-like behavior or locomotion in the open field (O, unpaired *t*-test: *t*_10_ = 0.11, *P* = 0.91; P, unpaired *t*-test: *t*_10_ = 0.69, *P* = 0.50) or on the elevated-plus maze (Q, unpaired *t*-test: *t*_10_ = 0.31, *P* = 0.76; R, unpaired *t*-test: *t*_10_ = 0.27, *P* = 0.79). Data points are individual mice with mean ± s.e.m. Scale bars: white = 50 *μ*m, yellow = 500 *μ*m. * *P* < 0.05; ** *P* < 0.01; *** *P* < 0.001.

**Figure S16.**
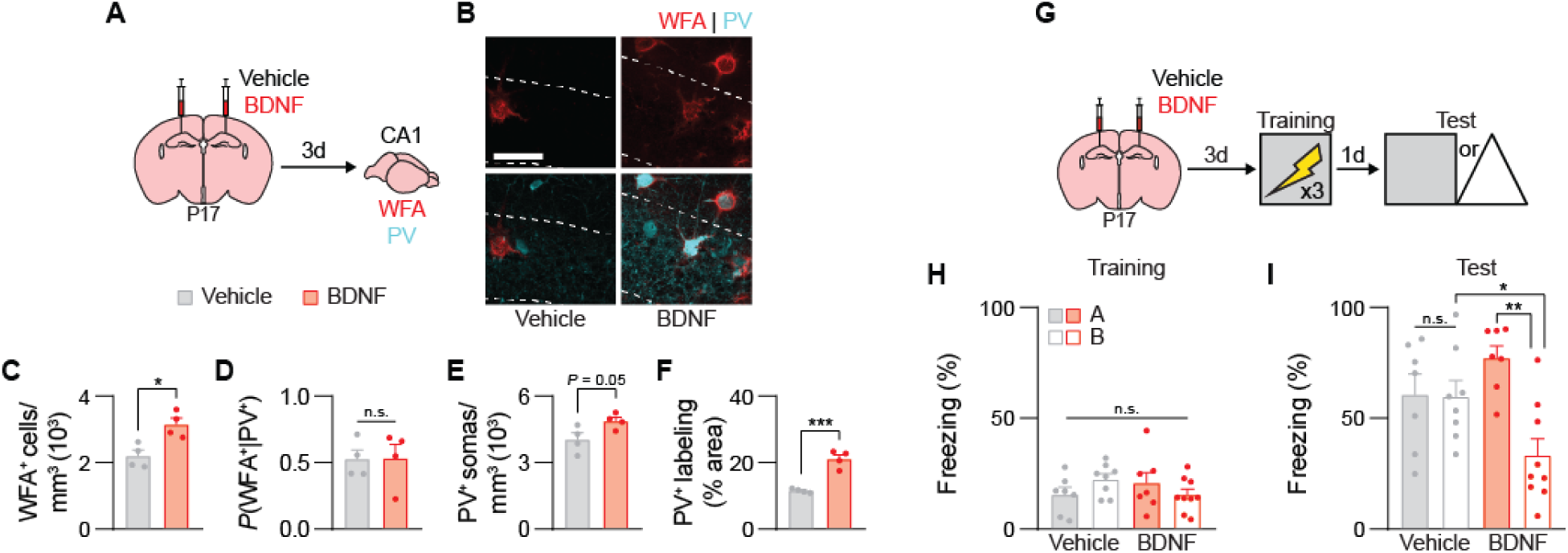
Accelerating CA1 PNN development with BDNF results in early onset of adult-like memory precision. **(A to B)** Recombinant BDNF protein was microinjected into the dorsal CA1 of juvenile mice (A) to promote PNN maturation (B). **(C to F)** BDNF treatment increased the density of WFA^+^ PNNs in the P20 CA1 (C, unpaired *t*-test: *t*_6_ = 3.64, *P* < 0.05). The proportion of PV^+^ interneurons surrounded by WFA^+^ PNNs was not increased (D, unpaired *t*-test: *t*_6_ = 0.051, *P* = 0.96), as BDNF also increased the number of PV^+^ interneurons (E, unpaired *t*-test: *t*_6_ = 2.42, *P* = 0.05) and density of PV^+^ neurites (F, unpaired *t*-test: *t*_6_ = 7.14, *P* < 0.001) in P20 CA1. **(G to I)** Juvenile mice were microinjected with Vehicle or BDNF 3 days before contextual fear conditioning (G). BDNF treatment did not alter freezing behavior during training (H, ANOVA, no Injection × Test Context interaction: *F*_1,27_ = 3.44, *P* = 0.074; no main effect of Injection: *F*_1,27_ = 0.07, *P* = 0.78; no main effect of Test Context: *F*_1,27_ = 0.07, *P* = 0.79) but improved memory precision during the test (I, ANOVA, Injection × Test Context interaction: *F*_1,27_ = 8.04, *P* < 0.01; no main effect of Injection: *F*_1,27_ = 0.43, *P* = 0.51; main effect of Test Context: *F*_1,27_ = 8.75, *P* < 0.01). Data points are individual mice with mean ± s.e.m. Scale bars: white = 50 *μ*m. * *P* < 0.05; ** *P* < 0.01.

